# Haplotype mining panel for genetic dissection and breeding in *Eucalyptus*

**DOI:** 10.1101/2022.08.23.503551

**Authors:** Julia Candotti, Nanette Christie, Raphael Ployet, Marja M. Mostert-O’Neill, S. Melissa Reynolds, Leandro Gomide Neves, Sanushka Naidoo, Eshchar Mizrachi, Tuan A. Duong, Alexander A. Myburg

## Abstract

To improve our understanding of genetic mechanisms underlying complex traits in plants, a comprehensive analysis of gene variants is required. *Eucalyptus* is an important forest plantation genus that is highly outbred. Trait dissection and molecular breeding in eucalypts currently relies on biallelic SNP markers. These markers fail to capture the large amount of haplotype diversity in these species and thus multi-allelic markers are required. We aimed to develop a gene-based haplotype mining panel for *Eucalyptus* species. We generated 17 999 oligonucleotide probe sets for targeted sequencing of selected regions of 6 293 genes implicated in growth and wood properties, pest and disease resistance and abiotic stress responses. We identified and phased 195 834 SNPs using a read-based phasing approach to reveal SNP-based haplotypes. A total of 8 915 target regions (at 4 637 gene loci) passed tests for Mendelian inheritance. We evaluated the haplotype panel in four *Eucalyptus* species (*E. grandis*, *E. urophylla*, *E. dunnii* and *E. nitens*) to determine its ability to capture diversity across eucalypt species. This revealed an average of 3.13 to 4.52 haplotypes per target region in each species and 33.36% of the identified haplotypes were shared by at least two species. This haplotype mining panel will enable the analysis of haplotype diversity within and between species and provide multi-allelic markers that can be used for genome-wide association studies and gene-based breeding approaches.

**Significance Statement:** We developed a haplotype sequencing panel for *Eucalyptus* targeting 8915 regions at 4637 gene loci associated with growth and wood properties, pest and disease resistance and abiotic stress response providing a genome-wide, multi-allelic, gene centric genotyping resource for eucalypts. We tested the panel in four *Eucalyptus* species (*E. grandis*, *E. dunnii*, *E. nitens* and *E. urophylla*) and found an average of 3.65 haplotypes per target region per species, and 9.98 across all four species.

## Introduction

Marker-trait associations are performed to improve our understanding of complex traits. The goal of such studies is to identify causative variants underlying phenotypes of interest and this information can be used in breeding programmes through marker-assisted breeding (MAB, Jiang, 2013). To perform genome-wide association analysis, a set of markers that sufficiently cover the genome is required. Biallelic single nucleotide polymorphisms (SNPs) are the most abundant source of polymorphic markers in plant genomes (Thudi *et al*., 2021) and can be detected using high-throughput methods such as SNP genotyping arrays (Silva-Junior *et al*., 2015) and sequencing-based genotyping (such as genotyping-by-sequencing (GBS, Deschamps et al., 2012)). Recently, there has been a shift towards multi-allelic haplotype-based (combinations of adjacent SNPs used as markers) association analysis in crop species such as rice (Ogawa, Yamamoto, *et al*., 2018; Ogawa, Nonoue, *et al*., 2018a), wheat (N’Diaye *et al*., 2017) and maize (Negro *et al*., 2019). Haplotype markers hold several advantages over SNPs, including increased polymorphic information content (N’Diaye *et al*., 2017), higher allelic diversity, and improved resolution in determining genomic positions of causal polymorphisms (Ogawa, Nonoue, *et al*., 2018b; Ogawa, Yamamoto, *et al*., 2018; Negro *et al*., 2019; Han *et al*., 2020). Furthermore, detection of interactions between haplotypes (epistasis) at different gene loci can explain some of the phenotypic variation of complex traits (Jan *et al*., 2019; Takeuchi *et al*., 2021). For highly heterozygous, outcrossing plants, multi-allelic haplotype markers are important to capture large amounts of genetic variation that cannot be identified using biallelic SNPs. For example, in a population constructed using two outcrossing individuals (Chen *et al*., 2021), there can be up to four allelic variants present, and biallelic SNPs cannot identify all four variants.

The two most common ways to identify SNP-based haplotype variants is based on a sliding window approach defined by a set number of SNPs, or based on linkage disequilibrium (LD) in overlapping segments (Lorenz *et al*., 2010). The SNP window method is challenging, as the optimal number of SNPs to include in a window is difficult to determine (Yang *et al*., 2006). The LD approach makes use of the observed LD to group adjacent SNPs, that are co-inherited, into haplotype blocks of variable length (Barrett *et al*., 2005). Despite the fact that LD varies across the genome, haplotype construction with this approach commonly use an average LD value (N’Diaye *et al*., 2017) and this can result in a decreased accuracy when defining haplotype blocks. Depending on the number of SNPs used and the LD decay, both of these approaches identify haplotypes that span multiple genes. While many studies have identified haplotypes using these two methods, typically by reanalysis of existing genome-wide SNP data (Bekele *et al*., 2018; Coffman *et al*., 2020; Jan *et al*., 2019), few studies have developed dedicated gene-based haplotype analysis tools.

Gene-based, multi-allelic haplotype markers allow gene-level resolution when performing genome-wide association studies which can enable the identification of causal variants within or near to genes of interest (Torkamaneh *et al*., 2021). Additionally, it is important to target cis-regulatory regions as these play an important role in quantitative trait variation (Wang *et al*., 2021). Gene-based haplotypes can subsequently be used for systems genetics, association analyses and functional genetics (Torkamaneh *et al*., 2021; Alonge *et al*., 2020). Genome-wide haplotype genotyping has been performed in rice (Yu *et al*., 2021; Zhang *et al*., 2021), soybean (Torkamaneh *et al*., 2021) and tomato (Alonge *et al*., 2020). These studies used resequencing data of 104 (Yu *et al*., 2021), 1007 (Torkamaneh *et al*., 2021) and 3024 (Zhang *et al*., 2021) accessions, respectively, to identify SNPs which were compiled into gene-centric haplotypes. However, obtaining genome sequencing data for a large number of individuals is not feasible or cost-effective in many plant species, leading to alternative approaches such as multiplexed, targeted resequencing to identify SNP-based haplotypes (Kamneva *et al*., 2017; Loera-Sánchez *et al*., 2021). There are a number of genomics service providers that enable custom targeted sequencing panel designs such as AmpliSeq (Illumina, San Diego, Callifornia, USA), QIAseq (Qiagen, Hilden, Germany) and Flex-Seq®Ex-L (Rapid Genomics, Gainesville, Florida, USA, referred hereafter as Flex-Seq).

*Eucalyptus* is a globally important tree genus, with over 700 recognized species (Ladiges *et al*., 2003). A number of fast-growing eucalypt species and their interspecific hybrids form the basis of a global hardwood fibre plantation industry (>20 mha world-wide, Iglesias and Wiltermann, 2009). Due to its economic importance, a number of genomic resources have been generated including an annotated reference genome (Myburg *et al*., 2014; Bartholomé *et al*., 2015), an Illumina EUChip60K SNP chip (Silva-Junior *et al*., 2015) and an Axiom 72K SNP chip (ThermoFisher Scientific, Waltham, Massachusetts, USA). These arrays, especially the EUChip60K chip, have been used extensively for association mapping (R. T. Resende *et al*., 2017; Mhoswa *et al*., 2020; Rafael Tassinari Resende *et al*., 2017) and genomic selection (Tan *et al*., 2017; Mphahlele *et al*., 2020; R. T. Resende *et al*., 2017). Ballesta et al., (2019) used an LD approach to extracted haplotype blocks from SNP data and subsequently used the haplotypes for genomic prediction in eucalypts. This study showed that the use of haplotypes resulted in improved predictive ability, especially for low-heritability traits, despite the fact that they could only extract 1 137 haplotype blocks from 14 422 informative SNPs. As the benefits of haplotype markers are increasingly being shown in crop species, such as *Brassica napus*(Jan *et al*., 2019), rice (Yu *et al*., 2021; Zhang *et al*., 2021), soybean (Torkamaneh *et al*., 2021), maize (Coffman *et al*., 2020; Mayer *et al*., 2020) and pigeonpea (Sinha *et al*., 2020), it is important to explore haplotype diversity in forest tree crops such as eucalypts. Forest trees have the added challenge of being highly outbred and harbouring large amounts of allelic variation, both of which can be addressed with more informative multi-allelic haplotype markers.

Here, we describe the development of a multi-species, gene-centric haplotype mining panel for commercially grown *Eucalyptus* trees. The study aimed to (i) prioritize 5 000 genes associated with growth and wood properties, pest and disease resistance, and abiotic stress response for targeted genome sequencing based on locus-specific probe sets (Flex-Seq, Rapid Genomics, Gainesville, FL), (ii) determine which probe sets produce informative haplotype data in four *Eucalyptus* species (*E. grandis, E. urophylla, E. dunnii* and *E. nitens*) as well as *E. urophylla* x *E. grandis* interspecific hybrids, and (iii) analyse haplotype diversity in the four species.

## Materials and methods

### Plant materials and DNA isolation

Twenty diverse individuals from each of four *Eucalyptus* species (*E. grandis, E. urophylla, E. nitens* and *E. dunnii*, **Supplemental Table S1**) from multiple provenances, were selected to ensure that the haplotype marker panel works across multiple *Eucalyptus* species. Additionally, 200 F_1_ hybrid individuals from 10 full-sib (FS) families of *E. grandis* x *E. urophylla* (**Supplemental Table S2**), together with the parents of these crosses, were selected to test the performance of the haplotype marker panel in interspecific hybrids and to perform tests for Mendelian inheritance of SNPs and haplotypes (**Supplemental Figure S1**). DNA was extracted from leaf or immature xylem tissue using the NucleoSpin®Plant II DNA extraction kit (Machery-Nagel, Germany). A total of 288 DNA samples were analyzed by Rapid Genomics LLC (Gainesville, Florida, USA) using the panel described below.

### Selection of candidate genes to target in the haplotype marker panel

A lines-of-evidence (LoE) approach was used to prioritize candidate genes to target in the haplotype panel. Published (**Supplemental Table S3**) and unpublished (mainly transcriptome) datasets were used to identify genes most likely to be involved in growth and wood traits, abiotic stress, and pest and disease resistance, as well as plastid and mitochondrial encoded genes. For the unpublished data, LoE were drawn from experiments that involved transcriptome experiments. Lines of evidence were assigned to each gene based on the number of datasets in which the gene was identified. The final selection of genes was made by selecting those with the highest number of lines of evidence in each category.

### Probe design by Rapid Genomics

Probe sets were designed by Rapid Genomics for the selected genes to target the following regions relative to the annotated transcription start site (TSS) and the 3’ end of the gene (Bartholomé *et al*., 2015), respectively, in windows of 0-500 bp, 500-1000 bp, 1000-1500 bp and 1500-2000 bp up- and downstream (**Supplemental Figure S2**), with each probe set targeting an average of 200 bp interval to be sequenced. Various combinations of probe sets were selected for each gene (**Supplementary File 1**) based on Flex-Seq probe set design criteria such as base pair composition (i.e. GC and homopolymer length), distance to target region, reduced chance of binding of the probes to non-target regions of the genome, and overall probe hybridisation kinetic metrics.

### SNP identification and quality control

Flex-Seq libraries were sequenced on Illumina NovaSeq S4 flow-cells with paired-end 150 cycles, generating an average of 1.61 million reads per sample. The first step in haplotype characterization was to identify SNPs for each target region (**Supplemental Figure S1**). Raw reads were demultiplexed into individual sample indexes, processed to remove residual adapter dimers and resulting short reads (Trimmomatic), followed by alignment of resulting reads to the *E. grandis* v2 reference genome using Burrows-Wheeler Aligner (BWA, Li & Durbin, 2009). BAM files were processed for SNP identification using Genome Analysis Toolkit (GATK, DePristo *et al*., 2011). Briefly, SNPs and indels were identified using HaplotypeCaller with the following settings; the output was an intermediate GVCF file (-ERC GVCF), the output contained all variants (--output-mode EMIT_ALL_CONFIDENT_SITES) and ploidy was set to 4n (-ploidy 4) to accommodate the possibility that a small proportion of probe sets would detect duplicated loci (i.e. up to four haplotypes). Next, the single-sample GVCFs generated were imported into a GenomicsDB datastore using GenomicsDBImport with the intervals .bed file representing the entire *E. grandis* v2 reference genome (Bartholomé *et al*., 2015). GATK’s GenotypeGVCFs tool as part of GATK was used to genotype the samples in the GenomicsDB. SNPs were selected using the SelectVariants tool and the VariantFiltration function was used to retain SNPs that had a quality by depth > 2 (QD < 2), variant confidence > 30 (QAUL < 30), strand bias (estimated by the symmetric odds ratio test) < 3 (SOR > 3.0), strand bias (estimated by using Fisher’s exact test) < 60 (FS > 60), mapping quality > 40 (MQ < 40), mapping quality > −12.5 (MQRankSum < −12.5), position of REF versus ALT alleles within reads > −8 (ReadPosRankSum < −8). Using BCFtools v1.12 (McKenna *et al*., 2010), biallelic SNPs with less than 20% missing data were retained.

### Modification of SNP genotypes using variant allele frequency

Since it was necessary to classify SNPs as tetraploid in the previous steps (to accommodate possible cases of probe binding to duplicated gene loci leading to up to four haplotypes in a single individual), heterozygous SNPs were confirmed using the ratio of reference to alternative allele calls (allelic balance) within individuals’ data. This was done upon observation that the allelic balance of some heterozygous calls were skewed (**Supplemental Figure S3**). First, the variant allele frequency (VAF) of high quality heterozygous SNPs was determined, using the FS family data. Genotypes were called using the same method as described in the above section, except with the ploidy set as 2n and SNPs and their VAF values for samples from seven FS families (with parental data available), separated by family, were analysed in SVS v8.7.1 (SVS, Golden Helix®, Inc. Bozeman, MT, USA). Homozygous SNPs were retained in the parents by selecting for SNPs with a minor allele frequency (MAF) < 0.01 and no missing data. Heterozygous SNPs in the parents (that were polymorphic in the F1 progeny) were retained by selecting for SNPs with a MAF = 0.5 and call rate > 0.8. Markers that violated expected Mendelian segregation within FS families were removed. The VAF data was filtered to only include SNPs that were heterozygous in all progeny. The VAF data from all FS families was merged, and the 5^th^ and 95^th^ percentiles of the VAF values were determined.

Second, a python script (https://github.com/joanam/scripts/blob/master/allelicBalance.py) was modified to edit the heterozygous SNP calls across the entire dataset based on their VAF values. Briefly, heterozygous SNPs were identified in the input file. If a heterozygous SNP had a VAF greater than 23% or less than 70% (5th and 95th percentiles identified in previous paragraph), the SNP was written to the output file as heterozygous 0/0/1/1. If the VAF was less than 23% or greater than 70%, a chi-square test was performed with an expected allele depth of 25% (tetraploid). If the SNP passed the chi-square test (p-value ≥ 0.05), the SNP was written to the output file as it was in the input file originally. If the SNP failed the chi-square test (p-value < 0.05), the genotype was converted to homozygous for the most common allele.

### Read-based phasing of SNPs and haplotype identification

To identify haplotypes at each of the target regions, a read-based phasing approach was undertaken using WhatsHap v1.1 (Martin et al., 2016, **Supplemental Figure S1**) This tool phases adjacent SNPs by identifying which alleles are present on the same reads. The input was the filtered SNPs in VCF format and the mapped reads in BAM format. The polyphase method was used with default settings. Following phasing, the intersect function of BEDTools v2.30.0 (Quinlan and Hall, 2010) was used to label SNPs within each target region. Since WhatsHap only phases SNPs if there are two or more heterozygous SNPs in a region, regions with single heterozygous SNP were manually assigned to two haplotypes. In cases where WhatsHap failed to phase two or more heterozygous SNPs, SNPs were flagged for downstream analyses.

### Haplotype quality control – Mendelian segregation of haplotypes in FS families

Mendelian segregation testing of haplotypes was performed in seven FS families (**Supplemental Table S2**) to identify high quality targets that produce haplotypes originating from a single genetic locus. Due to the fact that we anticipated some proportion of probe sets to bind to (unknown) gene duplicates and therefore called all SNPs using a tetraploid model, a small proportion of target regions had more than two haplotypes in some individuals (**Supplemental Figure S4**). These target regions, present within some individuals, were marked as missing data for Mendelian analysis. SVS v8.7.1 was used to perform a Mendelian error check with the number of Mendelian errors per marker recorded.

Target regions (probe sets) were classified into three quality categories based on their Mendelian segregation patterns of the resulting haplotypes in the seven FS families, parental haplotype call rate and haplotype call rate across FS families. Category 1 target regions passed the Mendelian check in all FS families (with allowance for one Mendelian error per FS family), had both parental haplotypes correctly called in at least one FS family, had >80% call rate in at least one FS family and had no unphased SNPs. Category 2 target regions, passed the Mendelian check in at least one FS family (with allowance for one Mendelian error per FS family) with no missing parent data and an 80% call rate in that FS family, but did not pass the Mendelian check in some FS families, or had unphased SNPs present in some of the other FS families. Category 3 target regions, did not pass the Mendelian check, had missing parental data, or had <80 call rate across all FS families.

To determine the percent heterozygosity for SNPs in Category 1 target regions, SNPs within the target regions were extracted using BCFtools v1.12 (McKenna *et al*., 2010) “view” command with a .bed file containing the positions of the target regions of interest. Individual heterozygosity was calculated by taking the number of heterozygous sites divided by the total number of SNPs called in that individual. To calculate the percent heterozygosity for the Category 1 haplotypes, the number of diploid, heterozygous haplotypes was divided by the total number of haplotypes called per individual.

### Haplotype quality control – Identification of target regions that contain more than two haplotypes per individual

Even though the probe sets were designed to target single copy sequences, some individuals may carry gene duplications that are not present in the V2.0 *E. grandis* reference assembly (Bartholomé *et al*., 2015) used for probe design. SNPs were called as tetraploid to enable the identification of off-target binding of probe sets in those individuals that may contain duplications of the target regions. This resulted in target regions containing more than two haplotypes in some individuals (**Supplemental Figure S4**). These regions were analysed to determine if they were due to known duplicated genes or due to off-target probe binding to an unknown sequence. The percentage of individuals carrying more than two haplotypes per target region was determined in the four species. The genes underlying these target regions were compared with known duplicated genes from the *E. grandis* v2.0 reference genome (Bartholomé *et al*., 2015). To test for enrichment of duplicated genes, we performed a chi-square test using the number of duplicated genes in the reference genome as the expected number of genes and the number of genes in the panel containing more than two haplotypes per individual as the observed number.

### Haplotype diversity analysis in the four *Eucalyptus* species

Haplotype diversity in the four *Eucalyptus* species was analysed using the Category 1 haplotypes. The number of haplotypes per target region was calculated within and across the four species as well as across the four target regions of each gene. Haplotype networks were generated for selected genes using pegas v1.1 (Paradis, 2010). Haplotype allele frequency was determined for all Category 1 haplotypes. Diploid SNPs (see section Modification of SNP genotypes using variant allele frequency) underlying Category 1 target regions were extracted using the “view”command of BCFtools v1.12 (McKenna *et al*., 2010), with a .bed file containing the positions of the target regions of interest. Minor allele frequency (MAF) of these SNPs, across all four species and one Half-sib (HS) family (**Supplemental Table S2**), was determined in SVS v8.7.1 (SVS, Golden Helix®, Inc. Bozeman, MT, USA).

### Gene ontology analysis

Gene Ontology (GO) biological process (GO-BP) enrichment was performed for all genes in the most (top 10%) and least (bottom 10%) diverse target regions to determine if specific gene classes were found in these two categories. GO-BPs terms were obtained per gene and functional enrichment and p-value correction for multiple testing were performed following the method described in Pinard *et al*., (2019). Enriched terms were selected if the p-value was less than 0.05.

### Reproducibility of SNP genotyping calls

The Flex-Seq®panel consisted of two groups of probe sets, with some overlap between the regions targeted, but no overlap in the probe sets. Each sample was analysed using both probe set groups. This enabled us to determine if the SNP calls were consistent in the overlap regions. Diploid SNPs were identified (see section **Modification of SNP genotypes using variant allele frequency)**in the data generated from the two groups of probe sets, with SNPs in each group being kept as separate .vcf files. SNPs that were found in both files were identified using BCFtools v1.12 (McKenna *et al*.,2010) “isec” function. The percentage of SNP calls that were identical across the two files was determined.

## Results

### Haplotype panel targets growth and wood property, pest and disease resistance and abiotic stress associated genes

To identify genes targeted in the Flex-Seq panel, a combination of published and in-house datasets were used as lines of evidence for gene selection (**Supplemental Table S3**). We aimed to target 5000 genes, but to account for potential limitations in probe design, a list of 7969 candidate genes were selected to represent growth and wood properties (5714 genes), pest and disease resistance (1732 genes), and abiotic stress responses (843 genes, **Supplemental Table S4**). A total of 6.40% of genes were represented in two or more categories (**Supplemental Figure S5**). The final probe set panel designed and produced by Rapid Genomics contained 17 999 probe sets targeting one or more regions of 6293 genes (**Supplementary File 1**). The number of genes in each category were 4253 genes for growth and wood properties, 1152 for pest and disease resistance and 504 for pest and disease resistance.

### Identification of high-quality SNPs

In order to identify haplotypes, individual SNPs were first called using the mapped sequencing reads obtained from Rapid Genomics. Following three SNP filtering steps, a total of 14 071 probe sets (target regions in 5 672 genes) containing 156 770 SNPs remained (**Supplemental Figure S6**). Despite avoiding duplicated sequences at the probe design stage (using the *E. grandis* V2.0 reference genome, Bartholomé et al., 2015), we still expected to recover some haplotypes from more than one target region in the genome. Therefore, to enable identification of target regions containing more than two haplotypes in some individuals, SNPs were called with the ploidy set as four (**Supplemental Figure S4**). Initial analysis of this data suggested that the SNP genotype identified sometimes did not match the observed variant allele frequency (VAF, **Supplemental Figure S3**). To address this, we determined the VAF distribution of 8 569 high-quality heterozygous SNPs in seven FS families. The distribution of VAF across all individuals in the seven FS families showed that the 5th percentile was 0.2347 and the 95th percentile was 0.7007 (**Supplemental Figure S7**). This information was used to adjust heterozygous SNP genotypes (see Materials and Methods).

### A genome-wide panel that captures SNP and haplotype diversity

We identified a total of 14 071 probe sets, targeting 5 672 genes, following SNP read-based phasing in WhatsHap v1.1 (Martin *et al*., 2016) with target regions distributed genome-wide (**Supplemental Figure S8**) except for chromosome 5 which exhibited a number of regions with low density and putative positions of centromeres on other chromosomes. Across all samples (individuals) analysed, we were able to call haplotypes for an average of 88.13% of the 14 071 target regions (**Supplemental Figure S9**).

To determine if the panel captured sufficient SNP variation to identify haplotype diversity, we analysed the number of SNPs and haplotypes per target region in the four species. The mean number of SNPs per target region was 11.14 (**Figure 1A**), equating to the possibility of detecting 2 048 haplotypes per target region. The mean number of haplotypes per target region was 11.22 (**Figure 1B**), indicating that there are more than sufficient numbers of SNPs per target region to detect the observed haplotype diversity (**Figure 1D**). The number of haplotypes per target region was proportional to the number of SNPs per target region (**Figure 1C**).

**Figure 1.**
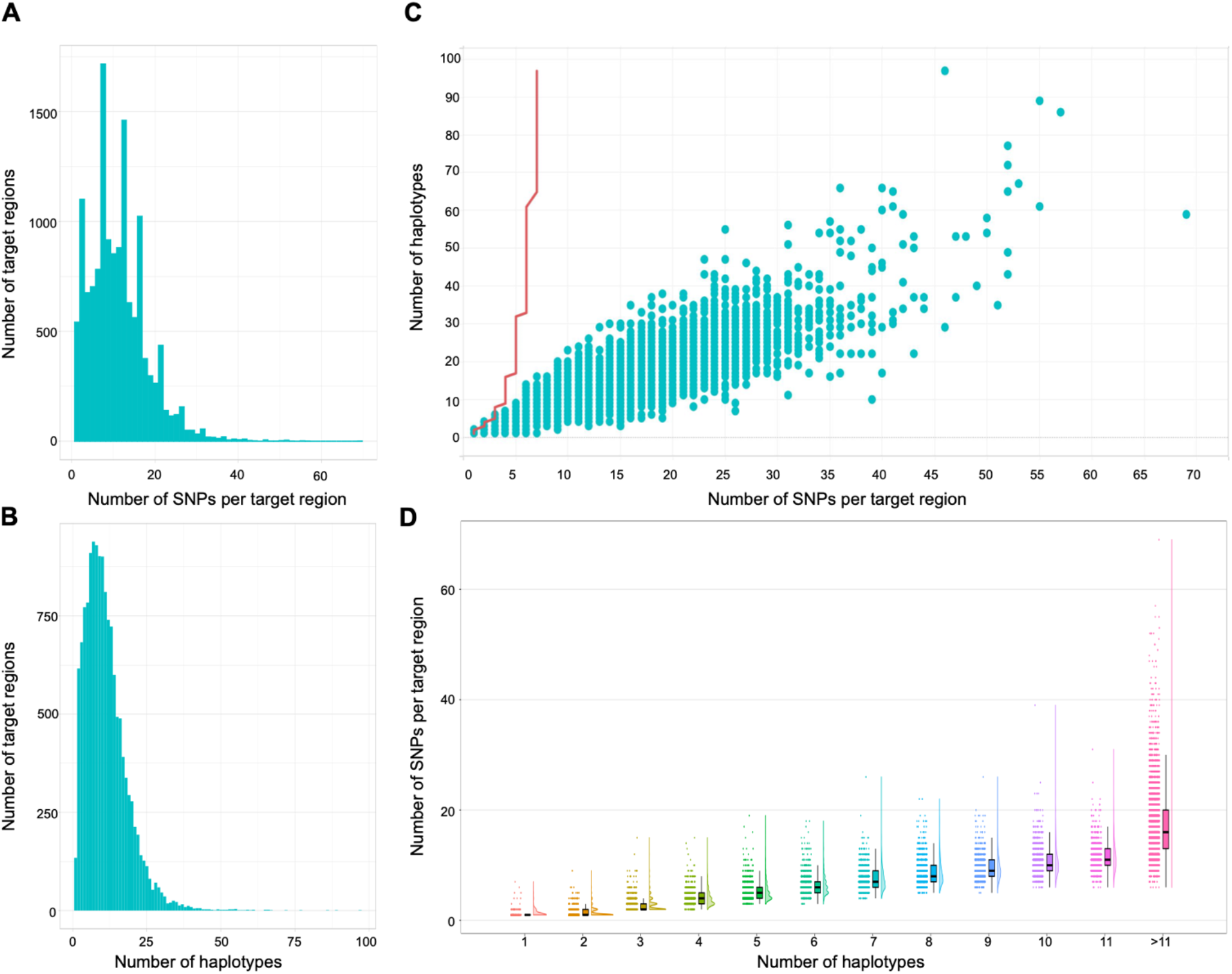
Genome-wide haplotype and SNP diversity captured by the haplotype marker panel. **A.**Distribution of the number of target regions with the given number of SNPs per target region (median = 10). **B.** Distribution of the number of target regions with a given number of haplotypes (median = 14). **C.** Number of observed haplotypes and corresponding SNPs per target region and the maximum number of haplotypes possible given the number of SNPs (red line). **D.** Distribution of the number of SNPs per target region for the given number of haplotypes.

Next, we used the segregation patterns of the haplotypes in seven FS families to identify high-quality haplotypes. We separated the haplotype blocks into three categories based on the number of Mendelian segregation errors, call rate and missing parent information across the seven FS families. Category 1 (high quality haplotypes) contained 8 915 target regions and 4 637 genes, Category 2 contained 4 227 target regions and 3 177 genes, and Category 3 contained 929 target regions and 844 genes (**Table 1**, see **Materials and Methods** for category definitions). We determined the physical positions of the target regions for each category (**Supplemental Figure S8**), and found that the target regions were found genome-wide. Category 1 target regions had significantly higher read depth compared to Category 2 and Category 3 target regions (**Supplemental Figure S10**).

**Table 1.** Summary of the number of target regions and haplotypes. Number of genes and number of haplotypes across the three target region categories, three gene categories and four species. “All” is the data across all four species combined.

| Gene category | Max number of haplotypes |  |  |  |  |  |  | Mean number of haplotypes |  |  |  |  |
| --- | --- | --- | --- | --- | --- | --- | --- | --- | --- | --- | --- | --- |
|  | No. target regions | No. genes | *E. grandis* | *E. urophylla* | *E. dunnii* | *E. nitens* | All | *E. grandis* | *E. urophylla* | *E. dunnii* | *E. nitens* | All |
| Category 1 haplotypes |  |  |  |  |  |  |  |  |  |  |  |  |
| Growth and wood properties | 6527 | 3348 | 22 | 21 | 20 | 21 | 72 | 3.70 | 4.52 | 3.29 | 3.12 | 9.94 |
| Pest and disease resistance | 1409 | 741 | 16 | 22 | 21 | 13 | 37 | 3.70 | 4.43 | 3.34 | 3.09 | 9.86 |
| Abiotic stress response | 546 | 316 | 26 | 24 | 18 | 20 | 60 | 4.14 | 4.86 | 3.84 | 3.47 | 11.08 |
| Category 2 haplotypes |  |  |  |  |  |  |  |  |  |  |  |  |
| Growth and wood properties | 3010 | 2255 | 20 | 29 | 19 | 21 | 53 | 4.71 | 5.39 | 3.87 | 3.72 | 12.35 |
| Pest and disease resistance | 712 | 538 | 16 | 20 | 19 | 16 | 42 | 4.59 | 5.32 | 3.86 | 3.63 | 12.00 |
| Abiotic stress response | 356 | 262 | 18 | 20 | 20 | 24 | 54 | 4.41 | 5.78 | 4.48 | 4.00 | 13.51 |
| Category 3 haplotypes |  |  |  |  |  |  |  |  |  |  |  |  |
| Growth and wood properties | 600 | 551 | 40 | 35 | 38 | 19 | 97 | 6.72 | 6.98 | 4.95 | 4.69 | 16.87 |
| Pest and disease resistance | 186 | 170 | 30 | 31 | 23 | 24 | 66 | 6.98 | 7.84 | 5.57 | 5.08 | 18.76 |
| Abiotic stress response | 109 | 92 | 33 | 29 | 20 | 25 | 77 | 8.37 | 8.13 | 6.32 | 5.71 | 20.13 |

For a low percentage of target regions we observed three or four haplotypes in some individuals. On average, 2.74% of target regions contained three haplotypes and 0.31% contained four haplotypes (per individual) across the 288 samples. To assess whether some of these target regions with more than two haplotypes could be the result of local duplication events, we evaluated the physical position and percentage of target regions with more than two haplotypes per individual (**Supplemental Figure S11**). We found that these target regions were distributed throughout the genome and there were indeed some loci with high frequency of putatively duplicated regions, some of which appeared to be species-specific. On average, *E. grandis* had the lowest proportion of individuals with target regions containing more than two haplotypes per individual (2.42%) while *E. urophylla* has the highest (3.52%, **Supplemental Table S5**).

Next, we compared known duplicated genes, identified using the *E. grandis* v2 reference genome (Bartholomé *et al*., 2015) with genes at target regions with more than two haplotypes per individual. This comparison was undertaken to determine if the presence of three or four haplotypes was due to known gene duplication events or non-specific probe binding (due to unknown duplicates). Target regions were selected for the duplication analysis if they contained three or more haplotypes in 5%, 10%, 15% and 20% of the 288 samples (**Supplemental Table S6**). We detected significantly fewer duplicates than expected at all percentages, compared to the genome-wide frequency of known duplicates (**Supplemental Table S6**) consistent with the design criteria used for the Flex-Seq assays.

We also analysed the heterozygosity of the SNPs and the haplotypes for Category 1 target regions for all 288 samples. A total of 89 231 SNPs and 8 915 haplotypes were analysed. We found that the mean SNP heterozygosity was 7.71% and the mean haplotype heterozygosity was 39.38% (**Supplemental Figure S12**). These results confirm that, as expected, the multi-allelic haplotype markers are more polymorphic than the underlying bi-allelic SNPs, which would be favourable for genetic dissection studies.

### A multi-species, gene-centric haplotype marker panel

We analysed the call rate and number of haplotypes of Category 1 (high quality) target regions to determine the performance of the haplotype panel across the four species. We found that *E. grandis* had the highest call rate (**Supplemental Table S7**), while *E. dunnii* had the lowest call rate. The mean number of haplotypes remained consistent (at approximately three to four haplotypes per target region) across the species (**Figure 2, Table 1**). We found there were both shared and unique haplotypes with *E. urophylla* having the highest number of unique haplotypes (18 551 haplotypes, **Supplemental Figure S13**). These results suggest that this method of haplotype identification performs consistently across different species and is able to detect haplotype diversity.

**Figure 2.**
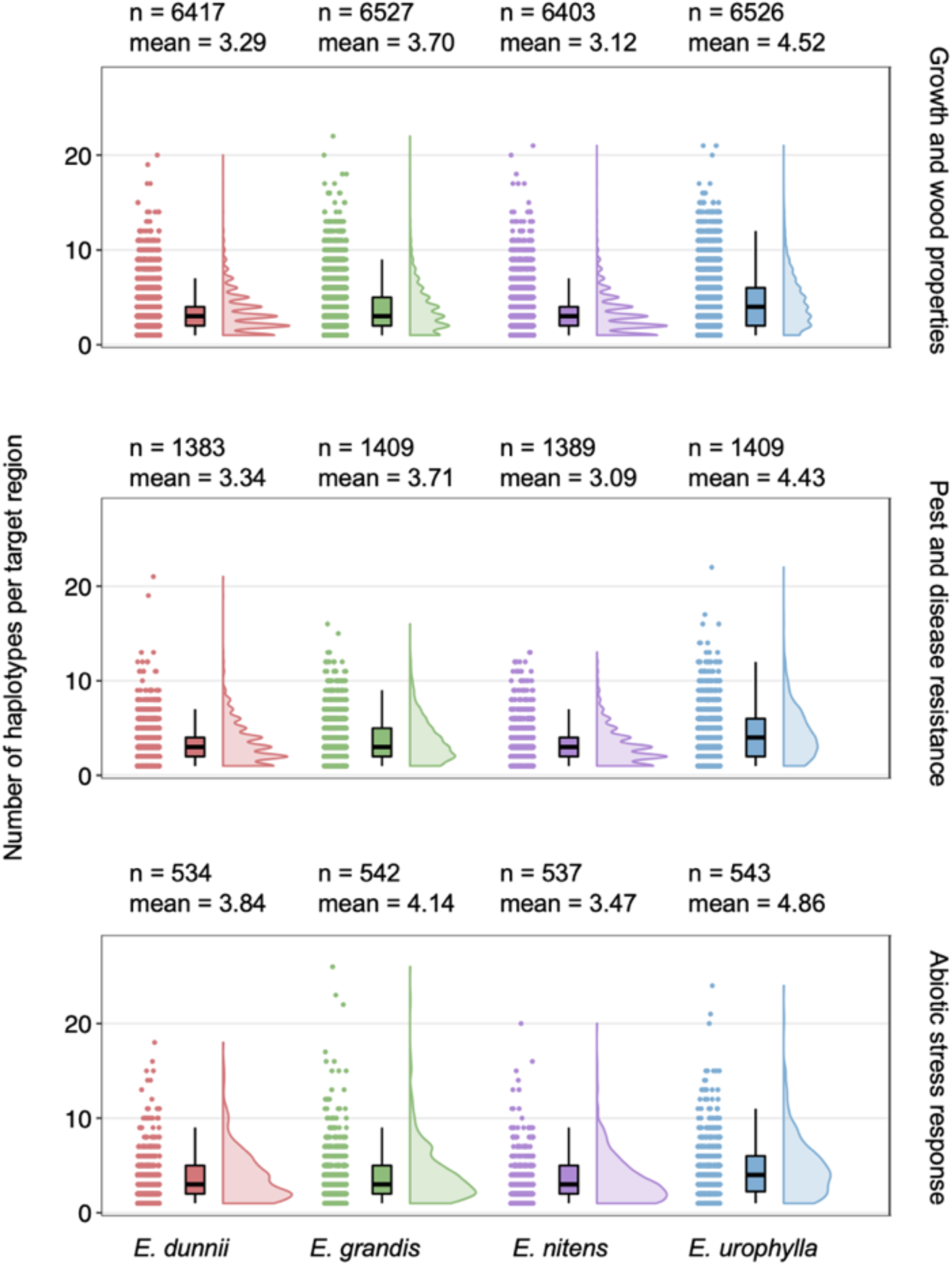
Haplotype diversity across gene categories in four *Eucalyptus* species. The number of haplotypes per target region (y-axis) as recorded for each of the four species (x-axis). A total of 20 individuals were analysed per species making the theoretical maximum number of haplotypes equal to 40 per target region. The mean value shown above each graph is the average number of haplotypes per target region and n is the number of target regions analysed in each category. A breakdown of haplotype diversity across the four species is provided in **Table 1.**

We subsequently analysed the haplotype diversity of Category 1 target regions across the three gene groups (growth and wood properties, pest and disease resistance and abiotic response genes) and different gene regions (upstream, gene start, gene end and downstream regions). Similar haplotype diversity was observed across the three gene groups, with growth and wood properties and pest and disease resistance genes having approximately 10 haplotypes per target region and abiotic stress response having 11 haplotypes per target region (**Table 1**). A similar pattern was observed when looking at the number of haplotypes across gene categories and gene regions (**Supplemental Figure S14**). The haplotype diversity was lower in the upstream and gene start regions than in the gene end and downstream regions (**Supplemental Figure S14**). No strong pair-wise correlations were observed between the different gene regions (**Supplemental Figure S15**).

We determined the SNP minor allele frequency and haplotype frequency for Category 1 target regions across the four species and within one HS family. Across the species, we found that 29.91% and 32.57% of SNPs and haplotypes, respectively, had allele frequencies less than 0.01 and 62.89% and 66.59% of SNPs and haplotypes, respectively, had allele frequencies less than 0.05 (**Supplemental Table S8**, **Supplemental Figure S16**). For the HS family, we found that 2.09% and 6.26% of SNPs, and 12.25% and 37.57% of haplotypes had frequencies less than 0.01 and 0.05 respectively. Next, we compared the SNP calls in regions which overlapped between the two groups of probe sets, to determine the reproducibility of SNP genotyping using the Flex-Seq technology. We analysed the SNP genotype calls across all 288 samples and found that the 982 SNPs analysed had an average allelic concordance of 95.33%. Of these, 67.82% (666 SNPs) had an allelic concordance of more than 99% and 87.78% (862) had an allelic concordance of 95% or more.

Next, we performed a GO enrichment analysis for genes within Category 1 haplotypes with the least haplotype diversity (bottom 10%, 836 genes) and the highest haplotype diversity (top 10%, 828 genes). GO-BP terms “determination of bilateral symmetry”, “meristem initiation” and “regulation of secondary cell wall biogenesis” were overrepresented in the least diverse haplotypes (**Supplemental Table S9**). No overrepresented GO was identified for the genes with the most diverse haplotypes (**Supplemental Table S9**).

Finally, we evaluated the use of the haplotype panel to understand gene variant diversity in biological pathways, focusing on the lignin biosynthetic pathway as an example (Carocha *et al*., 2015, **Supplemental Figure S17A**). First, we determined the number of individuals carrying haplotypes shared across all species, three species, two species and single species (**Supplemental Figure S17B**). We found that there were differences in the haplotype sharing across all target regions in the pathway, with some being mostly conserved and others containing more unique haplotypes. Next, we analysed haplotype sharing patterns between target regions of a single gene, Eucgr.I01134 (**Supplemental Figure S17C**). This gene was selected as it contained haplotype data for all four gene regions. We observed that different regions of the same gene could exhibit different patterns of unique and shared haplotypes, with the upstream and gene start regions being more conserved compared to the gene end and downstream regions.

## Discussion

A haplotype panel of 17 999 probe sets targeting 6 923 genes was designed and successfully used for genotyping, resulting in 195 834 high-quality SNPs in 14 071 target regions of 5 672 genes. Using Mendelian segregation of haplotypes in FS families, we identified 8 915 high quality target regions for 4 637 genes. We used the haplotype marker panel to identify 80 409 discrete haplotypes in 80 individuals of *E. grandis, E. nitens, E. urophylla* and *E. dunnii* (average of three to four haplotypes per target region).

Our aim was to develop a resource that can be used for haplotype-based association genetic studies in eucalypts. The genes were selected based on a LoE approach, but were distributed across the genome, making the panel useful for a genome-wide dissection using multi-allelic markers. We opted to test the panel across multiple eucalypt species and hybrids to determine transferability, and analysed multiple FS families to enable testing for Mendelian segregation and identification of high-quality SNPs and haplotypes. A total of 63.36% of the target regions produced haplotype markers with Mendelian segregation patterns. Our study was limited somewhat by the number of individuals per species (20) and we only analysed one hybrid combination. Additionally, the lack of high-quality reference genomes for other *Eucalyptus* species (besides *E. grandis*) precluded *in silico* prediction of the probe binding success in the three non-reference species (*E. urophylla, E. dunnii* and *E. nitens*).Although there was a fair expectation of sequence conservation in and near gene sequences, we had to rely on empirical testing to determine transferability to those species. Despite these limitations, we were able to identify 8 915 high quality haplotypes tagging 4 637 genes within and across the four species.

The four *Eucalyptus* species selected for this study (*E. grandis, E. urophylla, E. dunnii*, and *E. nitens*) are important for the global forestry industry as they are among the “big nine” most widely planted eucalypts (Harwood, 2011). Based on a collection of 20 diverse individuals per species, we found that *E. urophylla* contained the largest number of haplotypes (average of 4.52 haplotypes per target region) and the highest percentage (51.22%) of unique haplotypes. *E. urophylla* is found on seven islands of Indonesia (Pepe *et al*., 2004), with some evidence of natural hybridization on some islands (Payn *et al*., 2008) and is therefore thought to be more diverse than the other three species. The call rate of haplotypes was lower in *E. dunnii* and *E. nitens* individuals compared with *E. grandis* and *E. urophylla*. This is expected as the *E. grandis* reference genome (Myburg *et al*., 2014) was used for probe design. Furthermore, *E. grandis* and *E. urophylla* both belong to section *Latoangulatae*, while *E. dunnii* and *E. nitens* are part of the taxonomically more distant section *Maidenaria* (Brooker, 2000). Future iterations of this panel could make use of genome assemblies from all four species to improve probe design and transferability.

We designed probes to target multiple regions of each candidate gene. This was done to increase the likelihood that at least one target region per gene would be informative and to enable the analysis of haplotype diversity across the different gene regions. As the species-level LD decay in *Eucalyptus* is within four to six kilobases (Silva-Junior and Grattapaglia, 2015), future versions of this panel can retain the most informative probe set(s) per gene. Additionally, reducing the number of target regions to probe will allow multiplexed sequencing of larger numbers of samples per lane, which will reduce the cost of the haplotype analysis per individual, and allow haplotype genotyping of larger populations.

A technical challenge of this data was the presence of three and four haplotypes per individual per target for a small proportion (3.05%) of target regions, likely due to probes binding to unknown duplicated gene regions in those individuals. We used the *E. grandis* v2 genome reference (Bartholomé *et al*., 2015) during the probe design stage, however, pan-genome variation could result in duplications not considered in the probe design process. Future studies can include more genome sequences to help reduce off target binding. Even though the proportion of putative off target calls was low, it complicated the SNP and haplotype calling phase of the study. Unexpected duplications may be a feature of highly heterogenous genomes such as those of outbred eucalypts.

Despite only having 20 individuals per species, the haplotype panel was successfully used to sample haplotype diversity both within and among the four species. The mean number of haplotypes per target region was 3.13 to 4.52 haplotypes per species and 9.98 among the four species, of which, on average, 33.36% were shared between two or more species. These are similar to the number of haplotypes identified by Ballesta *et al*. (2019). In their study, the authors analysed 2 092 SNPs in 1 137 blocks (avg 1.8, range 2-12 SNPs per block) revealing a total of 3 279 haplotypes (avg 2.88 per block) segregating in 646 *E. globulus* individuals from a progeny trial of 62 full-sib and three half-sib families. This comparison is complicated by the fact that the authors had a much smaller number of markers per haplotype block. With over 60 families, the true number of haplotypes per block may be higher than three. Nevertheless, it is interesting that our study detected an average of 3.13 to 4.52 haplotypes per species, despite using 20 diverse individuals per species and having a sufficient number of SNPs to detect a much large number of haplotypes (avg 11.14 SNPs per block allows for a theoretical detection of up to 2 048 per target region).

Future work will include designing probes for a second version of the haplotype marker panel. Design criteria will include retaining at least two Category 1 target regions per gene, adding new probe sets for genes that did not have informative probe sets, and adding genes that were not included in the first version of the panel, but have sufficient lines of evidence to justify their inclusion. The objective would be to reach an optimal number of genes, target regions and sequencing depth that will allow multiplexing of a large number of samples to reduce the cost per sample to be competitive with existing SNP chip products for *Eucalyptus*, while providing a more informative, multi-allelic genotyping dataset. Ultimately, the Flex-Seq technology will allow users the option to target different numbers of genes and samples depending on the application.

The haplotype panel provides a resource which can be used in a number of ways (**Figure 3**). First, the haplotypes can be used as multi-allelic markers for genome-wide association studies (**Figure 3A**). Second, epistatic interactions between haplotypes can be analysed to identify favourable haplotype combinations (**Figure 3B**). This information can then be used for haplotype-based breeding in *Eucalyptus*. Third, the haplotype diversity within and across gene regions can be used to improve our understanding of gene evolution through the use of haplotype trees (**Figure 3C**), haplotype networks (**Figure 3D**), and LD across genes and gene regions (**Figure 3E**). Segregation patterns of the haplotypes within interspecific hybrid progeny (**Figure 3F**) can be used to advance our knowledge of hybrid compatibility and combining ability.

**Figure 3.**
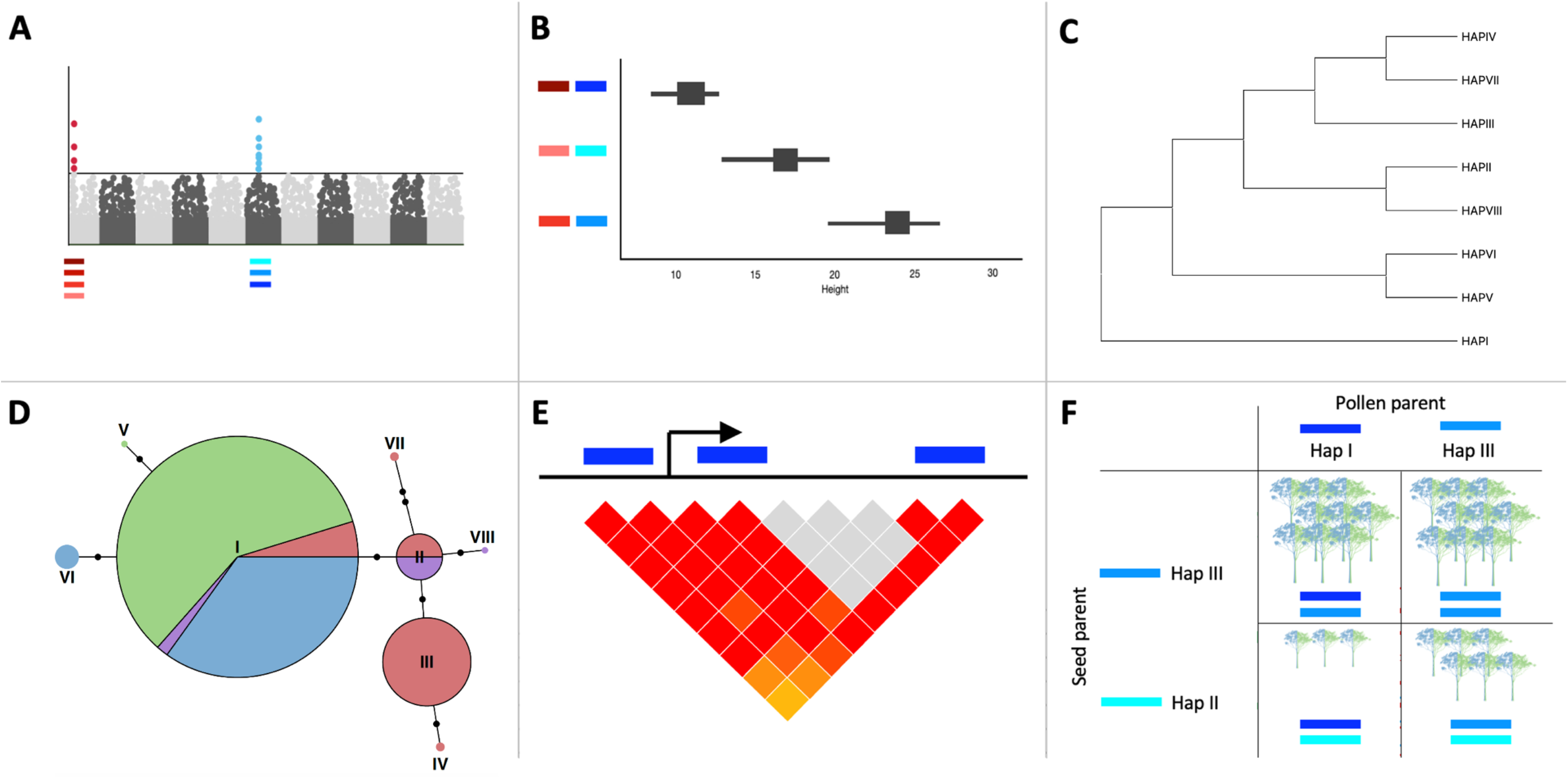
Overview of how the haplotype panel can be used in future studies. **A.** Haplotypes can be used as markers for genome-wide association studies. **B.** Epistatic interactions between haplotypes can be used to identify favourable haplotype combinations. Evolution of genes can be assessed through the use of (**C)** haplotype-based phylogenetic gene trees, **(D)** haplotype networks and **(E)** LD across gene regions (target regions shown in blue). **F.** Segregation patterns of haplotypes in interspecific hybrids can be used to analyse hybrid combining ability. Haplotypes are shown by coloured lines, the number of trees present represent the number of hybrid progeny carrying the specific haplotype combination. Haplotype combination Hap I/Hap II is present in the fewest individuals.

## Supplemental Material Statement

Supplemental tables and figures are provided as a separate document. There are nine Supplemental tables and 17 Supplemental figures. There is a one Supplementary excel file.

## Data Availability Statement

All of the short-read sequencing data are being submitted to the NCBI Short Read Archive (SRA) under the BioProject submission number SUB11917910.

## Author contributions

The idea was developed by AM. MO and SM assisted with sample selection and processing. JC, NC, RP, MO, SN performed gene selection. LGN supervised probe design. The data was analysed by JC with contributions from NC and TD and supervision from AM, EM, SN, TD. JC drafted the manuscript. All authors read, edited and approved by all authors.

## Acknowledgements

This work was funded in part by the Department of Science and Innovation and Technology Innovation Agency (DSI/TIA, Strategic Grant for the *Eucalyptus* Genomics Platform), the Forestry Sector Innovation Fund (FSIF *Eucalyptus* Genome Diversity Atlas grant), and by Sappi and Mondi through the Forest Molecular Genetics (FMG) Industry Consortium at the University of Pretoria. JC acknowledges PhD scholarship support from the NRF (UID 130369) and the University of Pretoria. We acknowledge the DNA Marker Team of the Forest Molecular Genetics Programme (FMG) for technical assistance. The authors thank the FMG Consortium members, in particular Sappi and Mondi, for genetic materials used in this study. Data processing was supported by the Centre for High Performance Computing (CHPC) in South Africa. The authors thank Anderson Silva (Rapid Genomics) for probe design and technical support.

## References

Alonge, M., Wang, X., Benoit, M., et al. (2020) Major impacts of widespread structural variation on gene expression and crop improvement in tomato. Cell, 182, 1–17. Available at: https://doi.org/10.1016/j.cell.2020.05.021.

Ballesta, P., Maldonado, C., Pérez-Rodríguez, P. and Mora, F. (2019) SNP and haplotype-based genomic selection of quantitative traits in *Eucalyptus globulus*. Plants, 8, 1–18. Available at: https://doi.org/10.3390/plants8090331.

Barrett, J.C., Fry, B., Maller, J. and Daly, M.J. (2005) Haploview: Analysis and visualization of LD and haplotype maps. Bioinformatics, 21, 263–265. Available at: https://doi.org/10.1093/bioinformatics/bth457.

Bartholomé, J., Mandrou, E., Mabiala, A., Jenkins, J., Nabihoudine, I., Klopp, C., Schmutz, J., Plomion, C. and Gion, J.M. (2015) High-resolution genetic maps of *Eucalyptus* improve *Eucalyptus grandis* genome assembly. New Phytol., 206, 1283–1296. Available at: https://doi.org/10.1111/nph.13150.

Bekele, W.A., Wight, C.P., Chao, S., Howarth, C.J. and Tinker, N.A. (2018) Haplotype-based genotyping-by-sequencing in oat genome research. Plant Biotechnol. J., 16, 1452–1463. Available at: https://doi.org/10.1111/pbi.12888.

Brooker, M. (2000) A new classification of the genus *Eucalyptus L’Hér*. (*Myrtaceae*). Aust. Syst. Bot., 13, 79–148.

Carocha, V., Soler, M., Hefer, C., Cassan-Wang, H., Fevereiro, P., Myburg, A.A., Paiva, J.A.P. and Grima-Pettenati, J. (2015) Genome-wide analysis of the lignin toolbox of *Eucalyptus grandis*. New Phytol., 206, 1297–1313. Available at: https://doi.org/10.1111/nph.13313.

Chen, M., Fan, W., Ji, F., et al. (2021) Genome-wide identification of agronomically important genes in outcrossing crops using OutcrossSeq. Mol. Plant, 14, 556–570. Available at: https://doi.org/10.1016/j.molp.2021.01.003.

Coffman, S.M., Hufford, M.B., Andorf, C.M. and Lübberstedt, T. (2020) Haplotype structure in commercial maize breeding programs in relation to key founder lines. Theor. Appl. Genet., 133, 547–561. Available at: https://doi.org/10.1007/s00122-019-03486-y.

DePristo, M.., Banks, E., Poplin, R., et al. (2011) A framework for variation discovery and genotyping using next-generation DNA sequencing data. Nat Genet., 43, 491–498. Available at: https://doi.org/10.1038/ng.806.A.

Deschamps, S., Llaca, V. and May, G.D. (2012) Genotyping-by-sequencing in plants. Biology (Basel)., 1, 460–483. Available at: https://doi.org/10.3390/biology1030460.

Han, Z., Hu, G., Liu, H., et al. (2020) Bin-based genome-wide association analyses improve power and resolution in QTL mapping and identify favorable alleles from multiple parents in a four-way MAGIC rice population. Theor. Appl. Genet., 133, 59–71. Available at: https://doi.org/10.1007/s00122-019-03440-y.

Harwood, C. (2011) New introductions – doing it right. In Developing a eucalypt resource: learning from Australia and elsewhere. pp. 43–54.

Iglesias, I. and Wiltermann, D. (2009) Eucalyptologics information resources on eucalypt cultivation.

Jan, H.U., Guan, M., Yao, M., et al. (2019) Genome-wide haplotype analysis improves trait predictions in *Brassica napus* hybrids. Plant Sci., 283, 157–164. Available at: https://doi.org/10.1016/j.plantsci.2019.02.007.

Jiang, G.-L. (2013) Molecular markers and marker-assisted breeding in plants. In Plant breeding from laboratories to fields. IntechOpen, pp. 45–83.

Kamneva, O.K., Syring, J., Liston, A. and Rosenberg, N.A. (2017) Evaluating allopolyploid origins in strawberries (*Fragaria*) using haplotypes generated from target capture sequencing. BMC Evol. Biol., 17, 1–19. Available at: https://doi.org/10.1186/s12862-017-1019-7.

Ladiges, P.Y., Udovicic, F. and Nelson, G. (2003) Australian biogeographical connections and the phylogeny of large genera in the plant family *Myrtaceae*. J. Biogeogr., 30, 989–998.

Li, H. and Durbin, R. (2009) Fast and accurate long-read alignment with Burrows-Wheeler transform. Bioinformatics, 25, 1754–1760. Available at: https://doi.org/10.1093/bioinformatics/btp698.

Loera-Sánchez, M., Studer, B. and Kölliker, R. (2021) A multispecies amplicon sequencing approach for genetic diversity assessments in grassland plant species. Mol. Ecol. Resour., 00, 1–21. Available at: https://doi.org/10.1111/1755-0998.13577.

Lorenz, A.J., Hamblin, M.T. and Jannink, J.L. (2010) Performance of single nucleotide polymorphisms versus haplotypes for genome-wide association analysis in barley. PLoS One, 5, 1–11. Available at: https://doi.org/10.1371/journal.pone.0014079.

Martin, M., Patterson, M., Garg, S., Fischer, S.O., Pisanti, N., Gunnar, W. and Marschall, T. (2016) WhatsHap: Fast and accurate read-based phasing. bioRxiv. Available at: https://doi.org/10.1101/085050.

Mayer, M., Hölker, A.C., González-Segovia, E., Bauer, E., Presterl, T., Ouzunova, M., Melchinger, A.E. and Schön, C.C. (2020) Discovery of beneficial haplotypes for complex traits in maize landraces. Nat. Commun., 11, 1–10. Available at: http://dx.doi.org/10.1038/s41467-020-18683-3.

McKenna, A., Hanna, M., Banks, E., et al. (2010) The genome analysis toolkit: A MapReduce framework for analyzing next-generation DNA sequencing data. Genome Res., 20, 1297–1303. Available at: https://doi.org/10.1101/gr.107524.110.

Mhoswa, L., O’Neill, M.M., Mphahlele, M.M., Oates, C.N., Payn, K.G., Slippers, B., Myburg, A.A. and Naidoo, S. (2020) A genome-wide association study for resistance to the insect pest *Leptocybe invasa* in *Eucalyptus grandis* reveals genomic regions and positional candidate defense genes. Plant Cell Physiol., 61, 1286–1296. Available at: https://doi.org/10.1093/PCP/PCAA057.

Mphahlele, M.M., Isik, F., Mostert-O’Neill, M.M., Reynolds, S.M., Hodge, G.R. and Myburg, A.A. (2020) Expected benefits of genomic selection for growth and wood quality traits in *Eucalyptus grandis*. Tree Genet. Genomes, 16, 1–12. Available at: https://doi.org/10.1007/s11295-020-01443-1.

Myburg, A.A., Grattapaglia, D., Tuskan, G.A., et al. (2014) The genome of *Eucalyptus grandis*. Nature, 510, 356–362. Available at: https://doi.org/10.1038/nature13308.

N’Diaye, A., Haile, J.K., Cory, A.T., Clarke, F.R., Clarke, J.M., Knox, R.E. and Pozniak, C.J. (2017) Single marker and haplotype-based association analysis of semolina and pasta colour in elite durum wheat breeding lines using a high-density consensus map. PLoS One, 12, 1–24. Available at: https://doi.org/10.1371/journal.pone.0170941.

Negro, S.S., Millet, E.J., Madur, D., Bauland, C., Combes, V., Welcker, C., Tardieu, F., Charcosset, A. and Nicolas, S.D. (2019) Genotyping-by-sequencing and SNP-arrays are complementary for detecting quantitative trait loci by tagging different haplotypes in association studies. BMC Plant Biol., 19, 1–22. Available at: https://doi.org/10.1186/s12870-019-1926-4.

Ogawa, D., Nonoue, Y., Tsunematsu, H., Kanno, N., Yamamoto, T. and Yonemaru, J.I. (2018a) Discovery of QTL alleles for grain shape in the Japan-MAGIC rice population using haplotype information. G3 Genes, Genomes, Genet., 8, 3559–3565. Available at: https://doi.org/10.1534/g3.118.200558.

Ogawa, D., Yamamoto, E., Ohtani, T., Kanno, N., Tsunematsu, H., Nonoue, Y., Yano, M., Yamamoto, T. and Yonemaru, J.I. (2018) Haplotype-based allele mining in the Japan-MAGIC rice population. Sci. Rep., 8, 1–11. Available at: https://doi.org/10.1038/s41598-018-22657-3.

Paradis, E. (2010) Pegas: An R package for population genetics with an integrated-modular approach. Bioinformatics, 26, 419–420. Available at: https://doi.org/10.1093/bioinformatics/btp696.

Payn, K.G., Dvorak, W.S., Janse, B.J.H. and Myburg, A.A. (2008) Microsatellite diversity and genetic structure of the commercially important tropical tree species *Eucalyptus urophylla*,endemic to seven islands in eastern Indonesia. Tree Genet. Genomes, 4, 519–530. Available at: https://doi.org/10.1007/s11295-007-0128-7.

Pepe, B., Surata, K., Suhartono, F., Sipayung, M., Purwanto, A. and Dvorak, W. (2004) Conservation status of natural populations of *Eucalyptus urophylla* in Indonesia & international efforts to protect dwindling gene pools. In Forest Genetic Resources. pp. 62–64.

Pinard, D., Fierro, A.C., Marchal, K., Myburg, A.A. and Mizrachi, E. (2019) Organellar carbon metabolism is coordinated with distinct developmental phases of secondary xylem. New Phytol., 222, 1832–1845. Available at: https://doi.org/10.1111/nph.15739.

Quinlan, A.R. and Hall, I.M. (2010) BEDTools: A flexible suite of utilities for comparing genomic features. Bioinformatics, 26, 841–842. Available at: https://doi.org/10.1093/bioinformatics/btq033.

Resende, R. T., Resende, M.D.V., Silva, F.F., Azevedo, C.F., Takahashi, E.K., Silva-Junior, O.B. and Grattapaglia, D. (2017) Assessing the expected response to genomic selection of individuals and families in *Eucalyptus* breeding with an additive-dominant model. Heredity (Edinb)., 119, 245–255. Available at: http://dx.doi.org/10.1038/hdy.2017.37.

Resende, Rafael Tassinari, Resende, M.D.V., Silva, F.F., Azevedo, C.F., Takahashi, E.K., Silva-Junior, O.B. and Grattapaglia, D. (2017) Regional heritability mapping and genome-wide association identify loci for complex growth, wood and disease resistance traits in *Eucalyptus*. New Phytol., 213, 1287–1300. Available at: https://doi.org/10.1111/nph.14266.

Silva-Junior, O.B., Faria, D.A. and Grattapaglia, D. (2015) A flexible multi-species genome-wide 60K SNP chip developed from pooled resequencing of 240 *Eucalyptus* tree genomes across 12 species. New Phytol., 206, 1527–1540. Available at: https://doi.org/10.1111/nph.13322.

Silva-Junior, O.B. and Grattapaglia, D. (2015) Genome-wide patterns of recombination, linkage disequilibrium and nucleotide diversity from pooled resequencing and single nucleotide polymorphism genotyping unlock the evolutionary history of *Eucalyptus grandis*. New Phytol., 208, 830–845. Available at: https://doi.org/10.1111/nph.13505.

Sinha, P., Singh, V.K., Saxena, R.K., et al. (2020) Superior haplotypes for haplotype-based breeding for drought tolerance in pigeonpea (*Cajanus cajan L.*). Plant Biotechnol. J., 1–9. Available at: https://doi.org/10.1111/pbi.13422.

Takeuchi, Y., Nishio, S., Terakami, S., Takada, N., Kato, H. and Saito, T. (2021) Haplotype structure analysis of a locus associated with fruit skin type on chromosome 8 in Japanese pear. Tree Genet. Genomes, 17, 1–13. Available at: https://doi.org/10.1007/s11295-020-01483-7.

Tan, B., Grattapaglia, D., Martins, G.S., Ferreira, K.Z., Sundberg, B. and Ingvarsson, P.K. (2017) Evaluating the accuracy of genomic prediction of growth and wood traits in two *Eucalyptus* species and their F1 hybrids. BMC Plant Biol., 17, 1–15. Available at: https://doi.org/10.1186/s12870-017-1059-6.

Thudi, M., Palakurthi, R., Schnable, J.C., et al. (2021) Genomic resources in plant breeding for sustainable agriculture. J. Plant Physiol., 257, 1–18. Available at: https://doi.org/10.1016/j.jplph.2020.153351.

Torkamaneh, D., Laroche, J., Valliyodan, B., et al. (2021) Soybean (*Glycine max*) Haplotype Map (GmHapMap): A universal resource for soybean translational and functional genomics. Plant Biotechnol. J., 19, 324–334. Available at: https://doi.org/10.1111/pbi.13466.

Wang, X., Aguirre, L., Rodríguez-Leal, D., Hendelman, A., Benoit, M. and Lippman, Z.B. (2021) Dissecting cis-regulatory control of quantitative trait variation in a plant stem cell circuit. Nat. Plants, 7, 419–427. Available at: http://dx.doi.org/10.1038/s41477-021-00898-x.

Yang, H.-C., Lin, C.-Y. and Fann, C.S.J. (2006) A sliding-window weighted linkage disequilibrium test. Genet. Epidemiol., 30, 531–545. Available at: https://doi.org/10.1002/gepi.

Yu, H., Li, Q., Li, Y., Yang, H., Lu, Z., Wu, J., Zhang, Z., Shahid, M.Q. and Liu, X. (2021) Genomics analyses reveal unique classification, population structure and novel allele of neo-tetraploid rice. Rice, 14, 1–16. Available at: https://doi.org/10.1186/s12284-021-00459-y.

Zhang, F., Wang, C., Li, M., et al. (2021) The landscape of gene–CDS–haplotype diversity in rice: Properties, population organization, footprints of domestication and breeding, and implications for genetic improvement. Mol. Plant, 14, 787–804. Available at: https://doi.org/10.1016/j.molp.2021.02.003.

