## Supplemental information for "Haplotype mining panel for genetic dissection and breeding in *Eucalyptus*"

29 pages, nine Supplemental Tables, 17 Supplemental Figures

### Supplementary Tables

#### Supplemental Table S1. Provenance and species information of 80 species samples.

| Sample ID | Species | Provenance |
|---|---|---|
| JN001 | *E. dunnii* | Lindesay Creek |
| JN002 | *E. dunnii* | Bald crab |
| JN003 | *E. dunnii* | Koreelah SF |
| JN004 | *E. dunnii* | Bald crab |
| JN005 | *E. dunnii* | Koreelah SF |
| JN011 | *E. dunnii* | Koreelah SF |
| JN014 | *E. dunnii* | Bald crab |
| JN016 | *E. dunnii* | Koreelah SF |
| JN020 | *E. dunnii* | Lindesay Creek |
| JN023 | *E. dunnii* | Lindesay Creek |
| JN026 | *E. dunnii* | Lindesay Creek |
| JN037 | *E. dunnii* | Oaky Hill |
| JZ034 | *E. dunnii* | Condamine |
| JZ048 | *E. dunnii* | Levuka |
| JZ055 | *E. dunnii* | Lindesay |
| JZ063 | *E. dunnii* | Acacia Creek |
| JZ066 | *E. dunnii* | Acacia Creek |
| JZ070 | *E. dunnii* | Dead horse track |
| JZ081 | *E. dunnii* | South Yabra |
| JZ089 | *E. dunnii* | South Yabra |
| 001_2 | *E. grandis* | Windsor Tablelands 1 |
| 014_2 | *E. grandis* | Windsor Tablelands 2 |
| 021_2 | *E. grandis* | Copperlode |
| 041_3 | *E. grandis* | Tinaroo Creek Road |
| 068_2 | *E. grandis* | Mt. Spec SF, Paluma |
| 080_3 | *E. grandis* | Paluma |
| 085_1 | *E. grandis* | Eungella |
| 091_2 | *E. grandis* | Finch Hatton Gorge |
| 100_2 | *E. grandis* | Brooweena SF |
| 128_3 | *E. grandis* | Yabba |
| 138_2 | *E. grandis* | Mapleton |
| 139_2 | *E. grandis* | Mapleton |
| 147_2 | *E. grandis* | Connondale |
| 157_3 | *E. grandis* | Mt. Mee |
| 167_2 | *E. grandis* | Mt. Tamborine |
| 172_3 | *E. grandis* | Mt. Lindesay |
| 199_2 | *E. grandis* | Bagawa SF |
| 201_2 | *E. grandis* | Wedding Bells SF |
| 213_3 | *E. grandis* | Orara West SF (72) |
| 233_2 | *E. grandis* | Bulahdelah SF |
| HG010 | *E. nitens* | Badja |
| HG020 | *E. nitens* | Badja |
| HG022 | *E. nitens* | Barrington tops |
| HG027 | *E. nitens* | Barrington tops |
| HG033 | *E. nitens* | Barrington tops |
| HG035 | *E. nitens* | Barrington tops |
| HG050 | *E. nitens* | Barren mountain |
| HG051 | *E. nitens* | Barren mountain |
| HG052 | *E. nitens* | Ebor |
| HG053 | *E. nitens* | Ebor |
| HG054 | *E. nitens* | Ebor |
| HG055 | *E. nitens* | Glenbog |
| HG056 | *E. nitens* | Glenbog |
| HG066 | *E. nitens* | New Zealand fri |
| HG078 | *E. nitens* | Tallaganda |
| HG079 | *E. nitens* | Tallaganda |
| HG082 | *E. nitens* | Landrace Kleinbuffelspruit* |

| | | |
|---|---|---|
| HG091 | *E. nitens* | Landrace Kleinbuffelspruit* |
| HG094 | *E. nitens* | Landrace Amsterdam van Aardt* |
| HG096 | *E. nitens* | Landrace Amsterdam van Aardt* |
| JW004 | *E. urophylla* | Dokeng |
| JW014 | *E. urophylla* | Kawela |
| JW035 | *E. urophylla* | Watololong |
| JW039 | *E. urophylla* | Apui |
| JW041 | *E. urophylla* | Mainang |
| JW067 | *E. urophylla* | Kilawair |
| JW075 | *E. urophylla* | Natakoli |
| JW080 | *E. urophylla* | Jontona |
| JW087 | *E. urophylla* | Padekluwa |
| JW113 | *E. urophylla* | Mollo |
| JW131 | *E. urophylla* | Tune |
| JW153 | *E. urophylla* | Nakana Ulam |
| JW155 | *E. urophylla* | Nesunhuhun |
| KN003 | *E. urophylla* | Jontona |
| KN012 | *E. urophylla* | Watakika |
| KN041 | *E. urophylla* | Koangao |
| KN052 | *E. urophylla* | Puor |
| KN055 | *E. urophylla* | Watakika |
| KN063 | *E. urophylla* | Koangao |
| KN070 | *E. urophylla* | Muda |

*Not F0 material

**Supplemental Table S2. Pedigree structure of the full-sib (FS) families used in this study.** The numbers represent the number of progeny per FS family with parentage confirmed

| | | *E. grandis* pollen parents | | |
|---|---|---|---|---|
| | | Parent 1 | Parent 2 | Parent 3[b] |
| *E. urophylla* seed parents | Parent 4 | 20[c,d] | 20[c] | 20 |
| | Parent 5 | 12[c,d] | 17[c] | 20 |
| | Parent 6[a] | 20 | | |
| | Parent 7[b] | 19[c,d] | | |
| | Parent 8 | 20[c,d] | | |
| | Parent 9 | 20[c,d] | | |

[a]No DNA was available for parent 6. Parent 6 is a known *E. grandis* seed parent.

[b]*E. urophylla* x *E. grandis* hybrid.

[c]FS families used for Mendelian segregation testing

[d]HS family used for minor allele frequency calculation

**Supplemental Table S3. Lines of evidence and gene numbers from published and unpublished data.** Both published data and in house datasets were used to create growth and wood property, pest and disease and environmental associated gene lists. Lines of evidence were created based on the study and the predicted function of the genes. Number of genes represents the total number of genes from each study/dataset that were added to the gene list.

| Study | Lines of evidence | Number of genes |
|---|---|---|
| Growth and wood property datasets | | |
| Myburg *et al.*, 2014 | 6 | 944 |
| Carocha *et al.*, 2015 | 1 | 39 |
| Mizrachi *et al.*, 2015 | 1 | 385 |
| Ployet *et al.*, 2019 | 3 | 3157 |
| Mizrachi *et al.*, 2017 | 4 | 1597 |
| Unpublished data | 40 | 31283 |
| Pest and disease datasets | | |
| Christie *et al.*, 2016 | 1 | 75 |
| Meyer *et al.*, 2016 | 2 | 161 |
| Mangwanda *et al.*, 2015 | 3 | 2140 |
| Oates *et al.*, 2015 | 3 | 1373 |
| Myburg *et al.*, 2014 | 1 | 134 |
| Tobias *et al.*, 2017 | 6 | 46 |
| Mhoswa *et al.*, 2020 | 1 | 15 |
| Külheim *et al.*, 2015 | 1 | 96 |
| Padovan *et al.*, 2015 | 2 | 27 |
| Zwart *et al.*, 2017 | 1 | 50 |
| Unpublished data | 13 | 36424 |
| Environmentally associated datasets | | |
| Mostert-O'Neill *et al.*, 2021 | 10 | 3643 |
| Ployet *et al.*, 2019 | 1 | 422 |
| Unpublished data | 6 | 7001 |

**Supplemental Table S4. Lines of evidence and number of genes for each category in the final gene list.**

| Category | Number of lines of evidence | Number of genes |
|---|---|---|
| Growth and wood properties | 55 | 5714 |
| Pest and disease resistance | 34 | 1732 |
| Abiotic stress response | 17 | 843 |
| Plastid genes | NA | 134 |
| Mitochondrial genes | NA | 67 |
| Total | NA | 7969* |

* Total are not calculated by adding the values together as there is overlap between categories

**Supplemental Table S5. Statistics for target regions with more than two haplotypes per individual for the four species.**

| Species | Number of target regions with more than two haplotypes in one or more individuals | Number of target regions with more than two haplotypes in more than 5% of individuals | Number of target regions with more than two haplotypes in more than 10% of individuals | Average percent of individuals with more than two haplotypes across target regions |
|---|---|---|---|---|
| *E. grandis* | 3260 (23.168%) | 3173 (22.55%) | 1254 (8.91%) | 2.42% |
| *E. urophylla* | 4307 (30.61%) | 3648 (25.93%) | 1663 (11.61%) | 3.52% |
| *E. nitens* | 2898 (20.60%) | 2094 (14.88%) | 1001 (7.11%) | 2.41% |
| *E. dunnii* | 2868 (20.38%) | 2685 (19.08%) | 1154 (8.20%) | 2.61% |
| All species | 7678 (54.57%) | 2358 (16.76%) | 775 (5.51%) | 2.72% |

**Supplemental Table S6. Number of target regions at genes known to be duplicated.** The number of genes containing three and four haplotypes in 5%, 10%, 15%, and 20% of 288 samples. The genes were compared to known duplicated genes annotated in Myburg *et al.* (2014). A chi-square test was performed to determine whether there was a significant difference between the observed and expected number of duplicated genes.

| Duplication class | Total number of genes in each duplication class based on v2 reference genome (percent of genes) | Observed | | | | Expected | p-value |
|---|---|---|---|---|---|---|---|
| | | Three alleles | Four alleles | Total | Percentage | | |
| | | | 5% | | | | |
| Tandem duplicates | 12560 (34.53%) | 546 | 93 | 639 | 11.27 | 1958.54 | <0.01 |
| Recent segmental duplicates | 279 (0.77%) | 1 | 0 | 1 | 0.02 | 43.67 | <0.01 |
| Lineage whole-genome duplication | 1877 (5.16%) | 211 | 10 | 221 | 3.90 | 292.68 | <0.01 |
| Ancient hexaploidization | 1788 (4.92%) | 208 | 7 | 215 | 3.79 | 279.06 | <0.01 |
| | | | 10% | | | | |
| Tandem duplicates | 12560 (34.53%) | 229 | 36 | 265 | 4.21 | 1958.54 | <0.01 |
| Recent segmental duplicates | 279 (0.77%) | 1 | 0 | 1 | 0.02 | 43.67 | <0.01 |
| Lineage whole-genome duplication | 1877 (5.16%) | 75 | 5 | 80 | 1.41 | 292.68 | <0.01 |
| Ancient hexaploidization | 1788 (4.92%) | 72 | 3 | 75 | 1.32 | 279.06 | <0.01 |
| | | | 15% | | | | |
| Tandem duplicates | 12560 (34.53%) | 123 | 20 | 143 | 2.27 | 1958.54 | <0.01 |
| Recent segmental duplicates | 279 (0.77%) | 0 | 0 | 0 | 0.00 | 43.67 | <0.01 |
| Lineage whole-genome duplication | 1877 (5.16%) | 36 | 4 | 40 | 0.71 | 292.68 | <0.01 |
| Ancient hexaploidization | 1788 (4.92%) | 28 | 1 | 29 | 0.51 | 279.06 | <0.01 |
| | | | 20% | | | | |
| Tandem duplicates | 12560 (34.53%) | 80 | 12 | 92 | 1.46 | 1958.54 | <0.01 |
| Recent segmental duplicates | 279 (0.77%) | 0 | 0 | 0 | 0.00 | 43.67 | <0.01 |
| Lineage whole-genome duplication | 1877 (5.16%) | 20 | 3 | 23 | 0.41 | 292.68 | <0.01 |
| Ancient hexaploidization | 1788 (4.92%) | 15 | 1 | 16 | 0.28 | 279.06 | <0.01 |

**Supplemental Table S7. Call rate of individuals and target regions for the four species across the three target region categories.** Prior to calculating the call rate, individual haplotype calls that contain three or four haplotypes were converted to missing.

| Call rate | *E. grandis* | *E. urophylla* | *E. nitens* | *E. dunnii* |
|---|---|---|---|---|
| Individuals with call rate above 80% | 18 (90%) | 17 (85%) | 19 (95%) | 13 (65%) |
| **Target region call rate** | | **All** | | |
| 0% | 2 (0.01%) | 4 (0.03%) | 378 (2.59%) | 328 (2.33%) |
| 80% | 12540 (89.12%) | 10582 (75.2%) | 10246 (73.82%) | 7096 (50.43%) |
| | | **Category 1 target regions** | | |
| 0% | 0 (0.00%) | 1 (0.01%) | 171 (1.92%) | 159 (1.78%) |
| 80% | 8367 (93.85%) | 7503 (84.16%) | 7086 (79.48%) | 5266 (59.07%) |
| | | **Category 2 target regions** | | |
| 0% | 0 (0.00%) | 0 (0.00%) | 175 (4.14%) | 142 (3.36%) |
| 80% | 3621 (85.66%) | 2720 (64.35%) | 2671 (63.19%) | 1572 (37.19%) |
| | | **Category 3 target regions** | | |
| 0% | 2 (0.22%) | 3 (0.32%) | 32 (3.44%) | 27 (2.91%) |
| 80% | 552 (59.42%) | 359 (38.64%) | 489 (52.64%) | 258 (27.77%) |

**Supplemental Table S8. Comparison of low frequency SNPs and haplotypes across the four species and a single half-sib family.** Prior to the analysis, target regions in individuals with more than two haplotypes were marked as missing.

| Allele frequency | Percent of SNPs | Percent of haplotypes |
|---|---|---|
| | Four Species | |
| <0.01 | 29.91 | 32.57 |
| <0.03 | 54.00 | 57.78 |
| <0.05 | 62.89 | 66.59 |
| | Half-sib family | |
| <0.01 | 2.09 | 12.25 |
| <0.03 | 3.08 | 28.47 |
| <0.05 | 6.26 | 37.58 |

**Supplemental Table S9. GO enrichment for the genes with the least (bottom 10%) diverse target regions.** Enrichment based on a chi-square test with a p-value threshold < 0.05.

| Term ID | Description | Genes in category | Cluster genes in category | Adjusted p-value |
|---|---|---|---|---|
| GO:0009855 | Determination of bilateral symmetry | 177 | 16 | 0.009 |
| GO:0010014 | Meristem initiation | 188 | 16 | 0.019 |
| GO:2000652 | Regulation of secondary cell wall biogenesis | 22 | 6 | 0.020 |

### Supplemental Figures

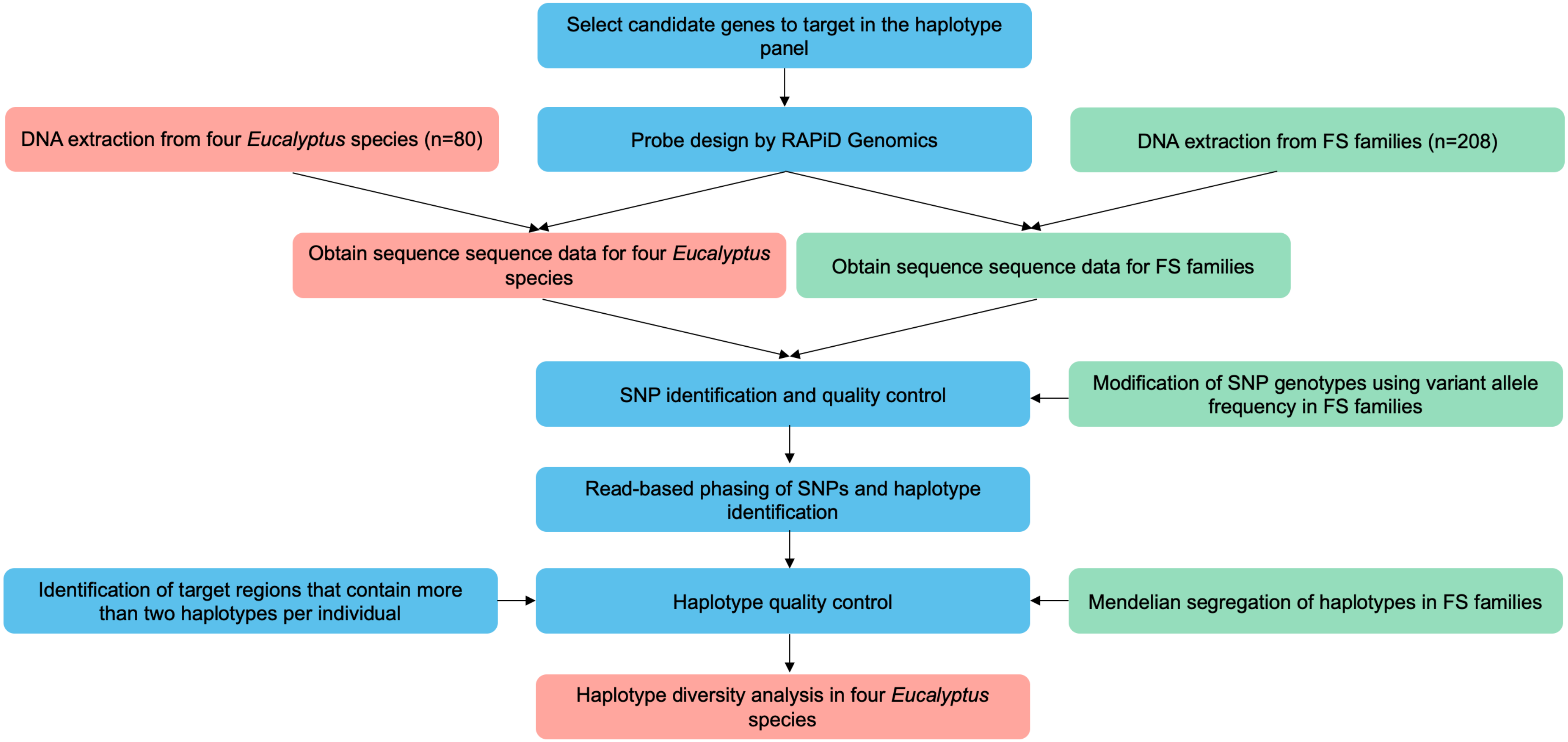


**Supplemental Figure S1. Overview of panel design process.** Blue indicates the step was performed on all samples/data, green indicates that the step was performed on Full-Sib (FS) family data/samples and red indicates the step was performed on the four *Eucalyptus* species data/samples. Grey represents the Gene Ontology (GO) enrichment analysis.

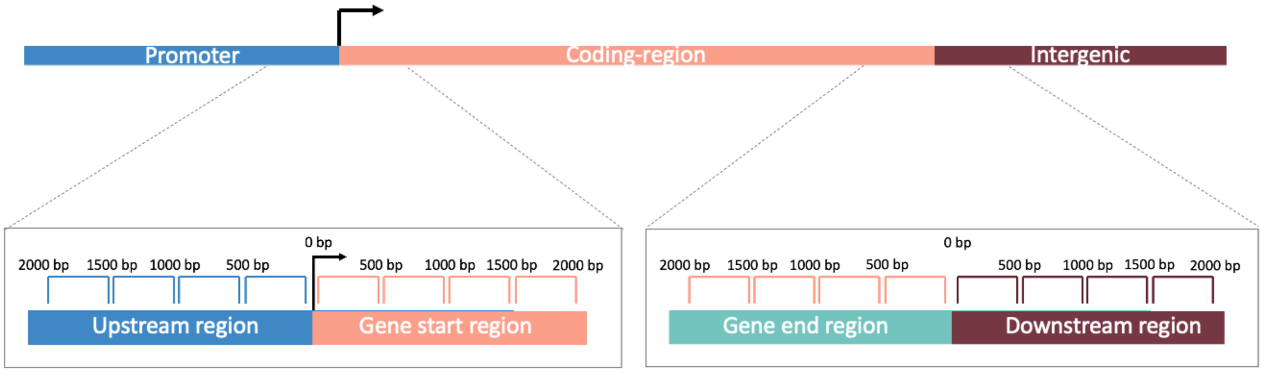


**Supplemental Figure S2. Target regions for probe design for each gene in the haplotype mining panel.** The black arrow marks the position of the annotated TSS.

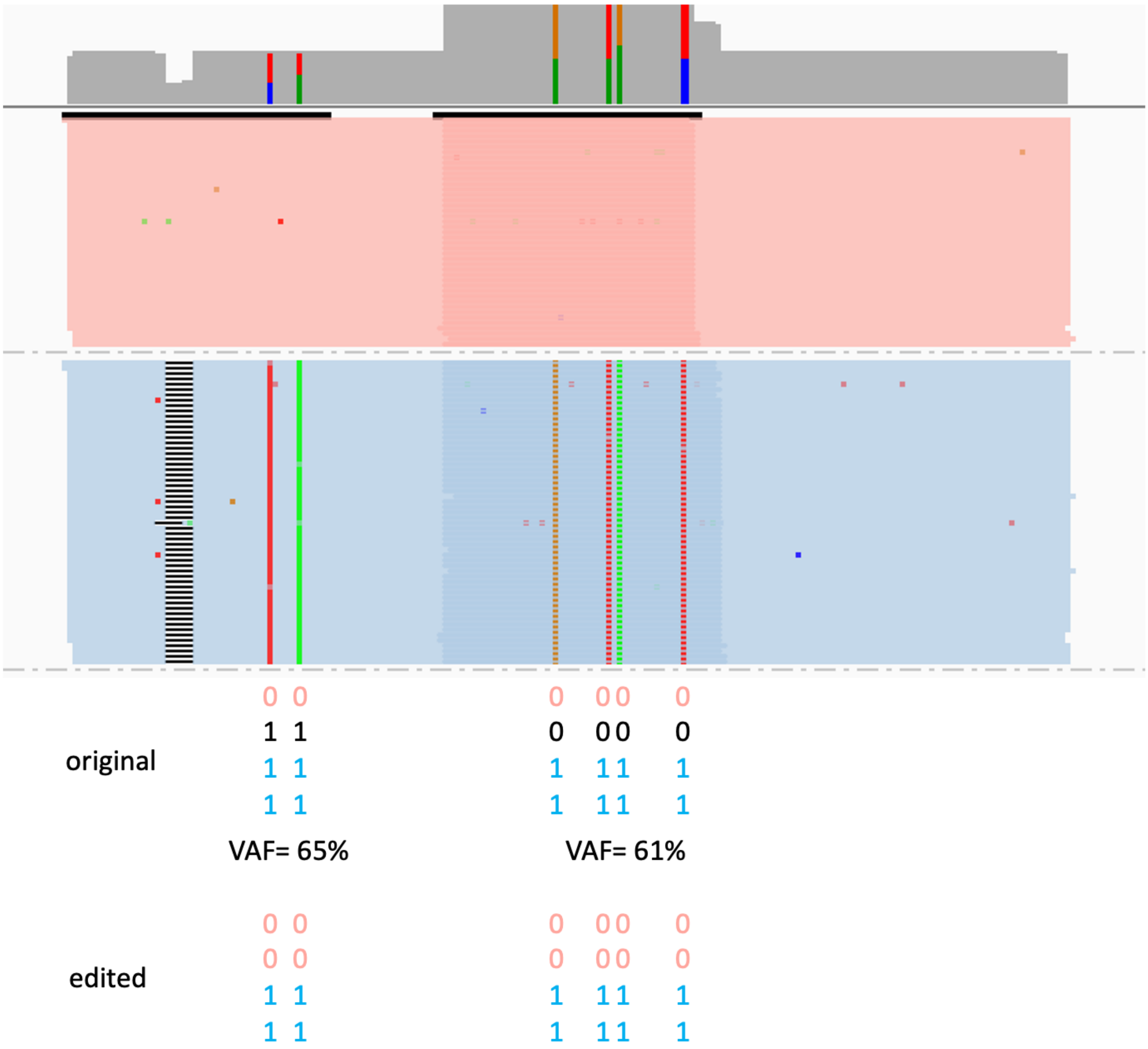


**Supplemental Figure S3. Example where the allelic balance of tetraploid SNPs caused incorrect haplotypes to be identified.** Reads (orange and blue) and coverage (grey) are shown in the top panel. Based on the reads, two haplotypes are present. When the SNPs are called as tetraploid, three haplotypes are identified by WhatsHap (Martin et al., 2016) in the original data using polyphase mode. However, the black haplotype is misidentified in this approach. Based on the variant allele frequency (VAF), we therefore correct the SNPs genotypes (see Materials and Methods). This resulted in two haplotypes being identified in the SNP data for this target region.

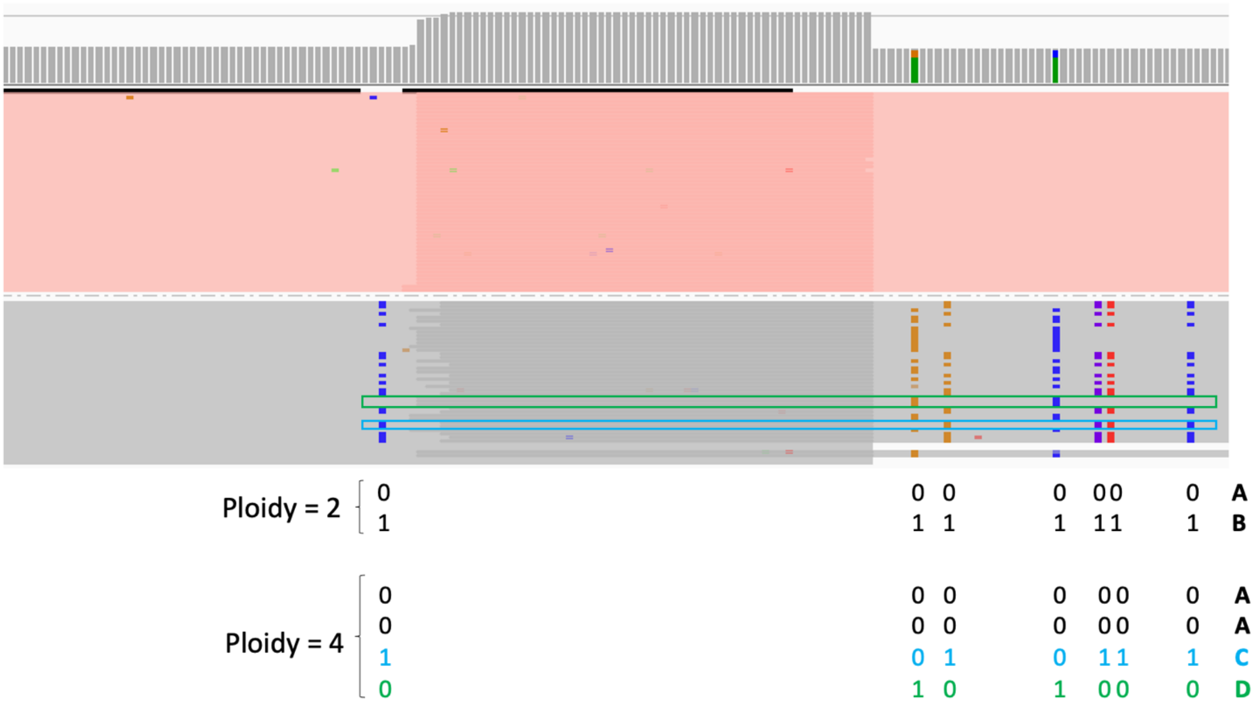


**Supplemental Figure S4. Example where calling SNPs as diploid lead to the identification of false haplotypes.** An overview of read depth is shown at the top (grey), the combinations of SNPs to create haplotypes are shown at the bottom (pink and grey reads). When SNPs are called as diploid, two haplotypes are identified, A and B. The A haplotype is a true haplotype (in the pink reads) but the B haplotype is false (combination does not exist). When SNPs are called as tetraploid, three real haplotypes are identified. Haplotype A is shown in the pink reads, an example of haplotype C is shown in the blue block and an example haplotype D is shown in the green block.

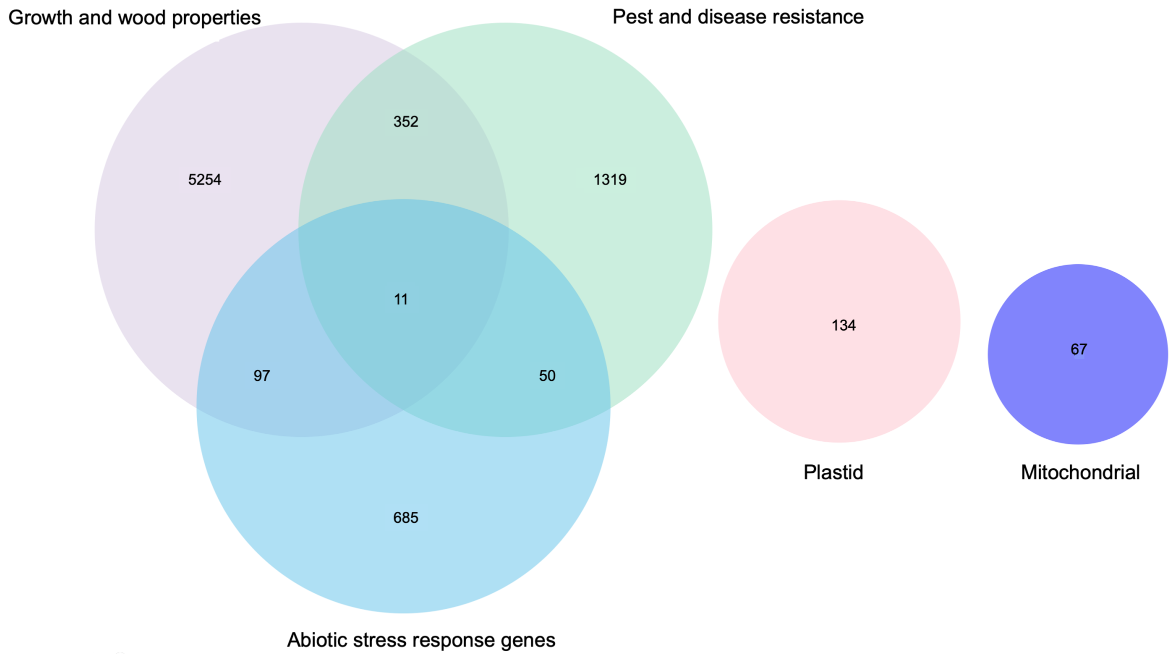


**Supplemental Figure S5. Number of genes in categories.** Number of genes in the main categories of growth and wood properties, pest and disease resistance, abiotic stress response, mitochondrial and plastid.

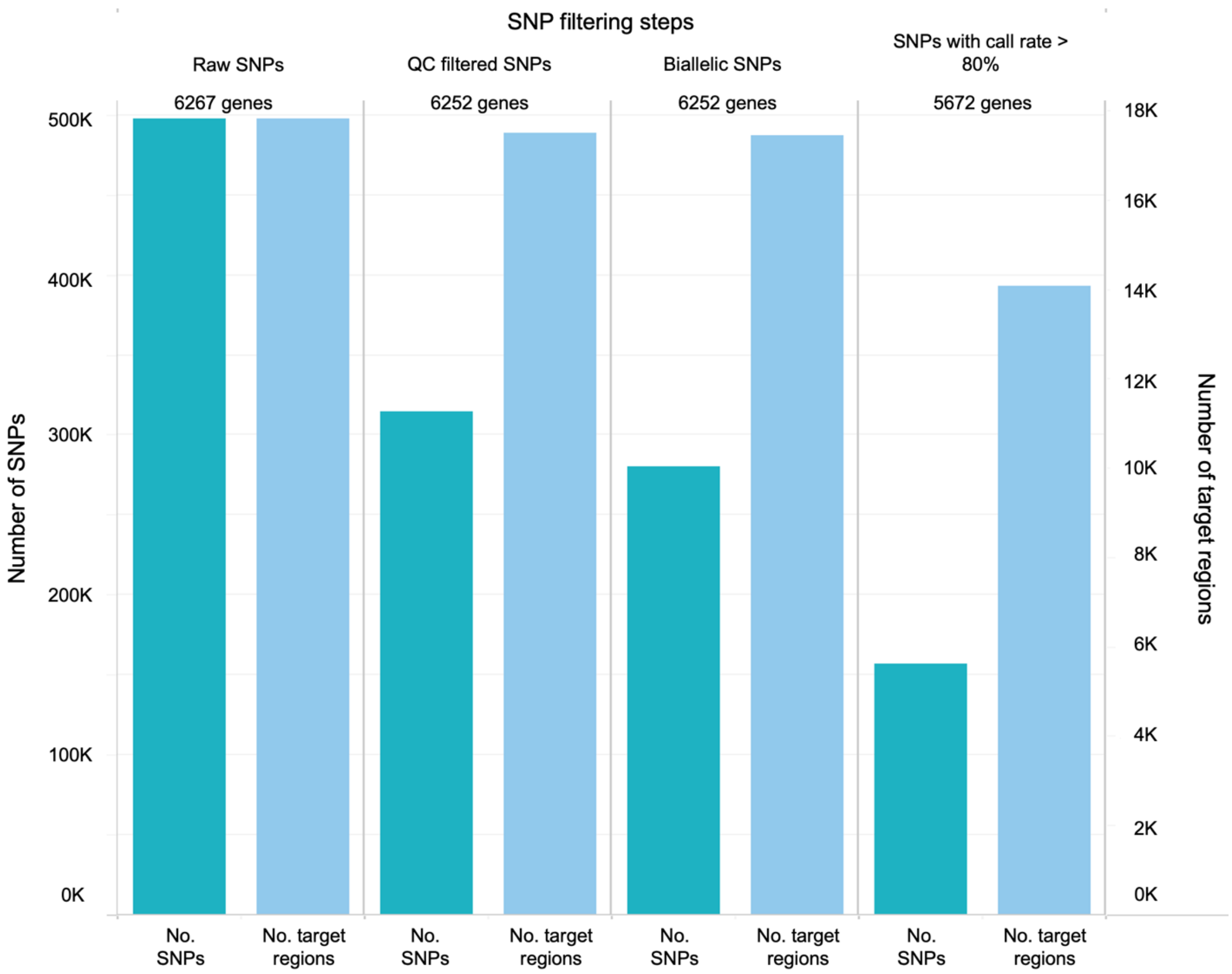


**Supplemental Figure S6. Number of SNPs and target regions remaining after each round of SNP filtering.** The y-axis on the left shows the number of SNPs and the y-axis on the right shows the number of target regions. Each filtering step is shown as a column, with the number of SNPs and number of haplotypes retained shown as bars and the number of genes retained as text.

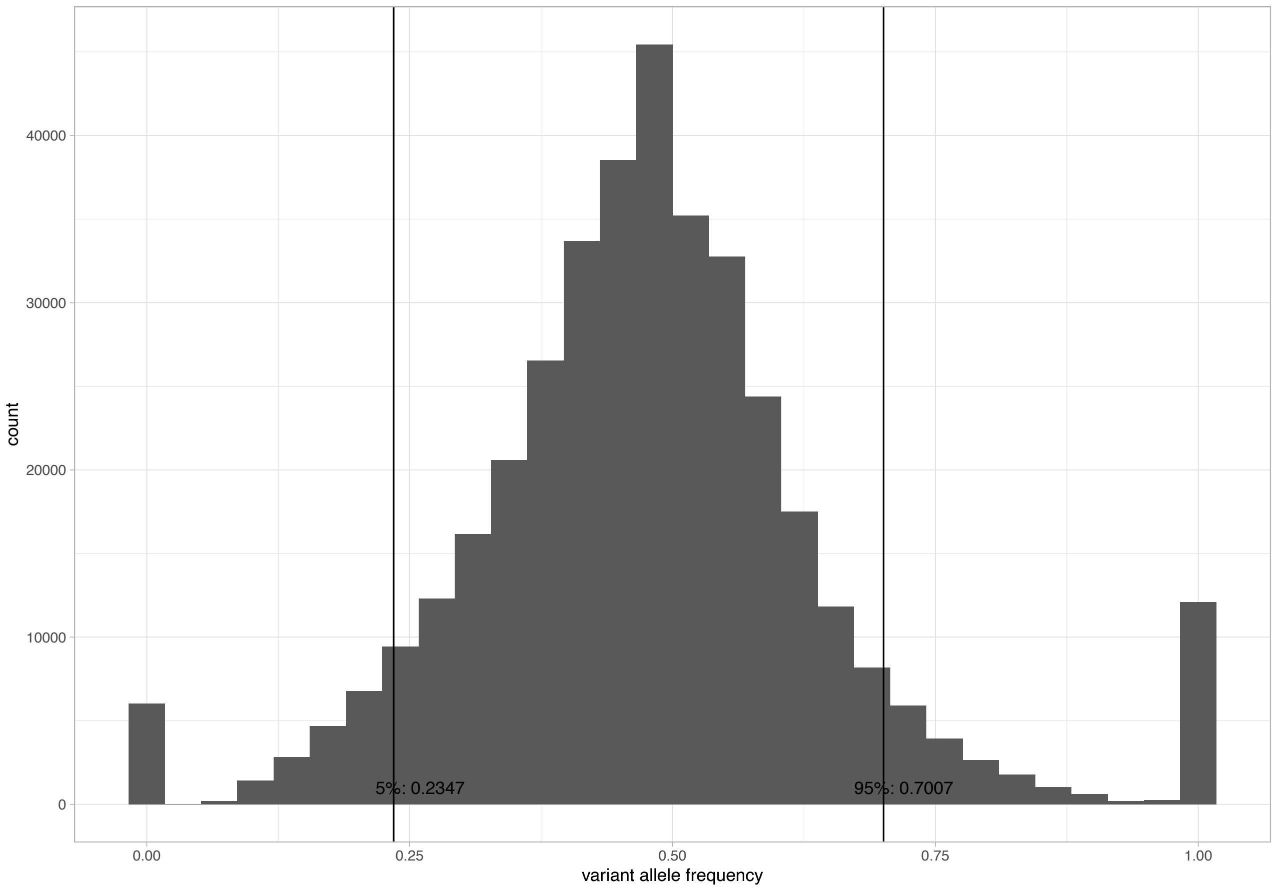


**Supplemental Figure S7. Distribution of SNP variant allele frequency across seven FS families.** A total of 8569 SNPs were identified as heterozygous in the seven FS families. The SNPs at 0 and 1 represent the homozygous parents. The 5th and 95th percentiles were calculated without the parents.

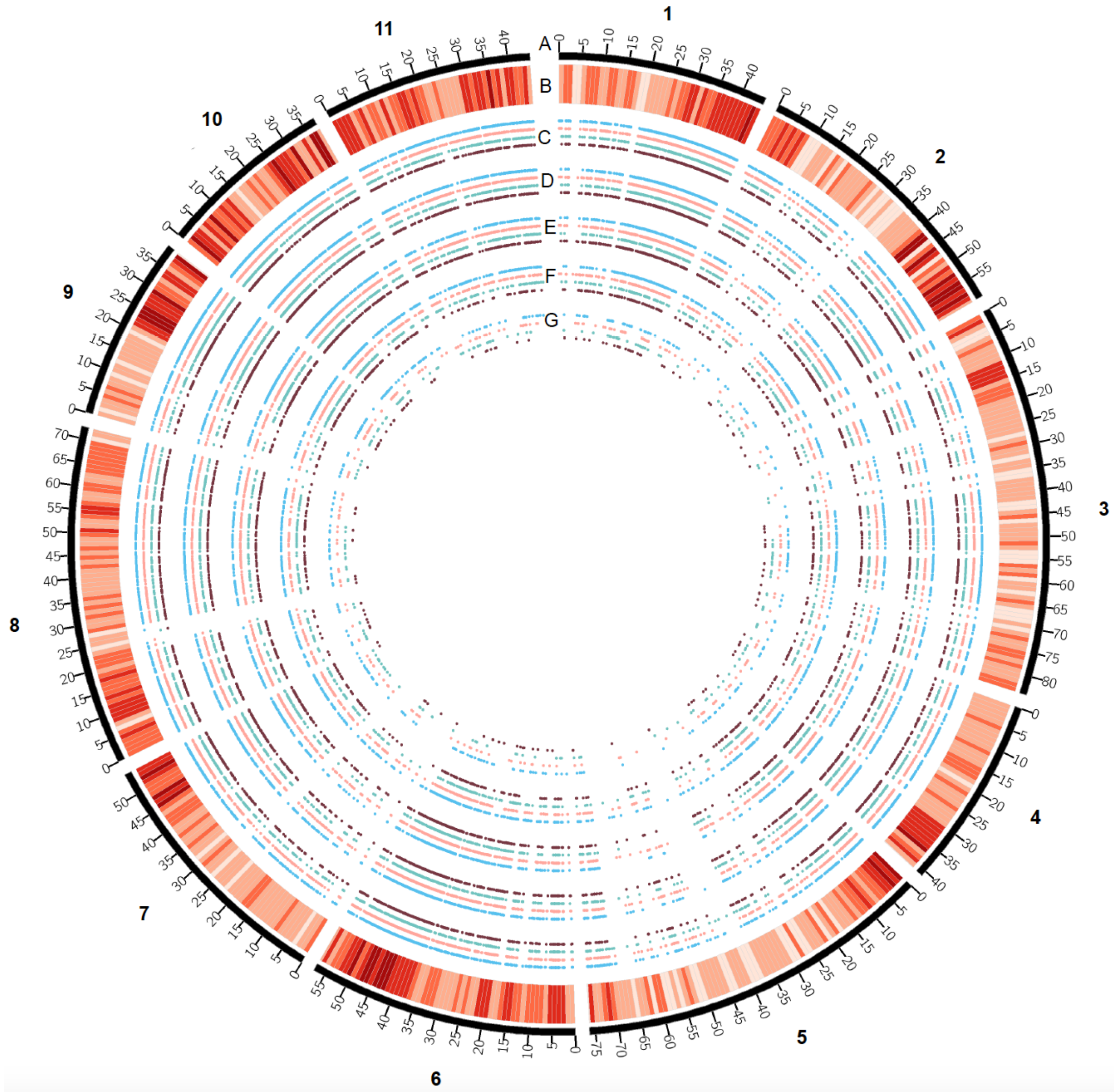


**Supplemental Figure S8. Location of target regions (haplotype markers) per haplotype quality category across the genome. A.** Megabase values and chromosomes based on the *E. grandis* v2 reference genome (Bartholomé et al., 2015). **B.** Gene density (number per Mb). **C.** Position of 17 999 probe sets. **D.** Position of 14 071 probe sets with SNPs. **E.** Position of 8 915 Category 1 target regions **F.** Position of 4 227 Category 2 target regions. **G.** Position of 929 Category 3 target regions. blue=upstream region, pink=gene start region, green=gene end region, black=downstream region.

A

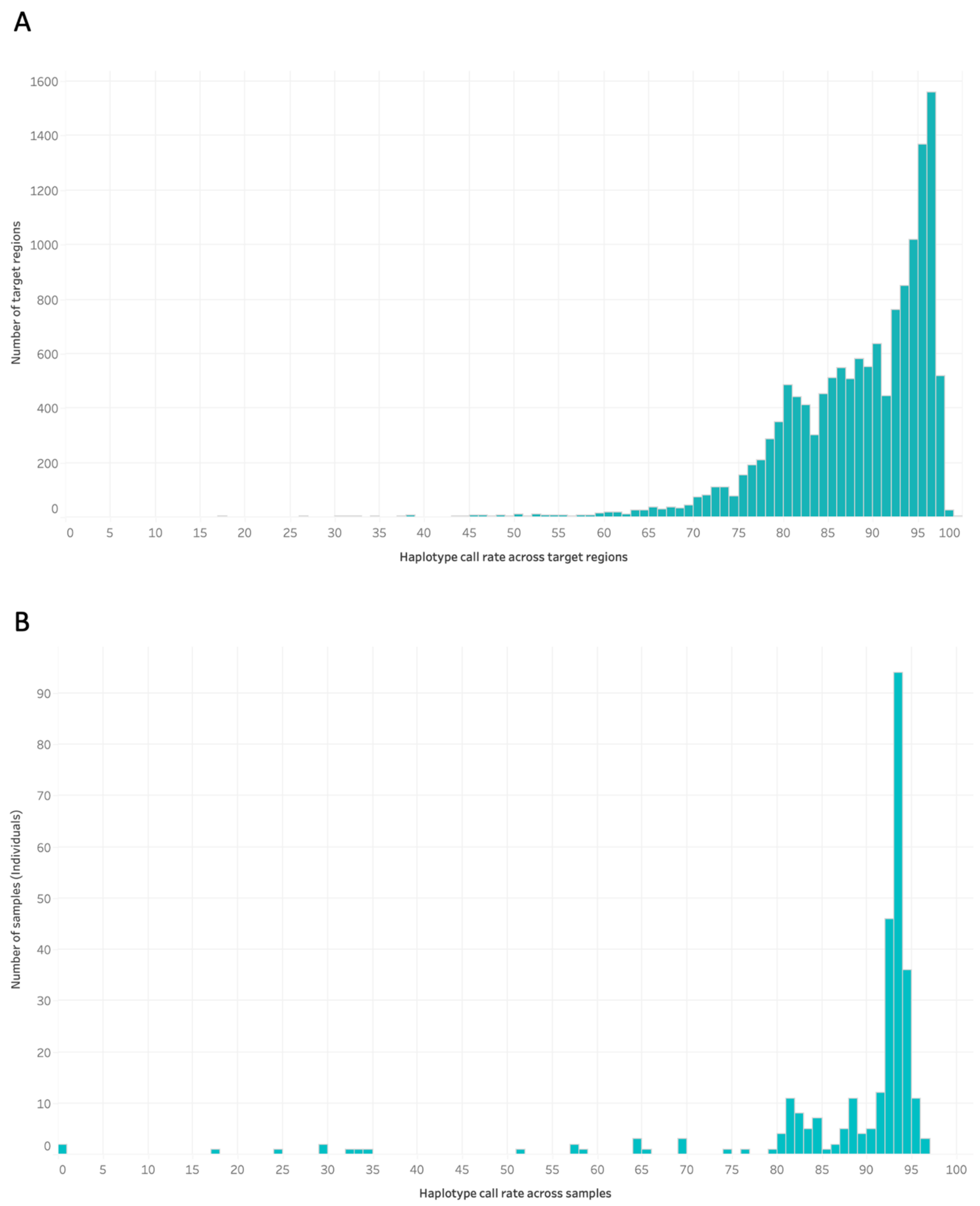


**Supplemental Figure S9. Distribution of haplotype call rate.** Prior to call rate being calculated, all haplotype calls with three or four haplotypes per individual were marked as missing **A.** Distribution of the haplotype call rate across target regions. **B.** Distribution of haplotype call rate across 288 samples.

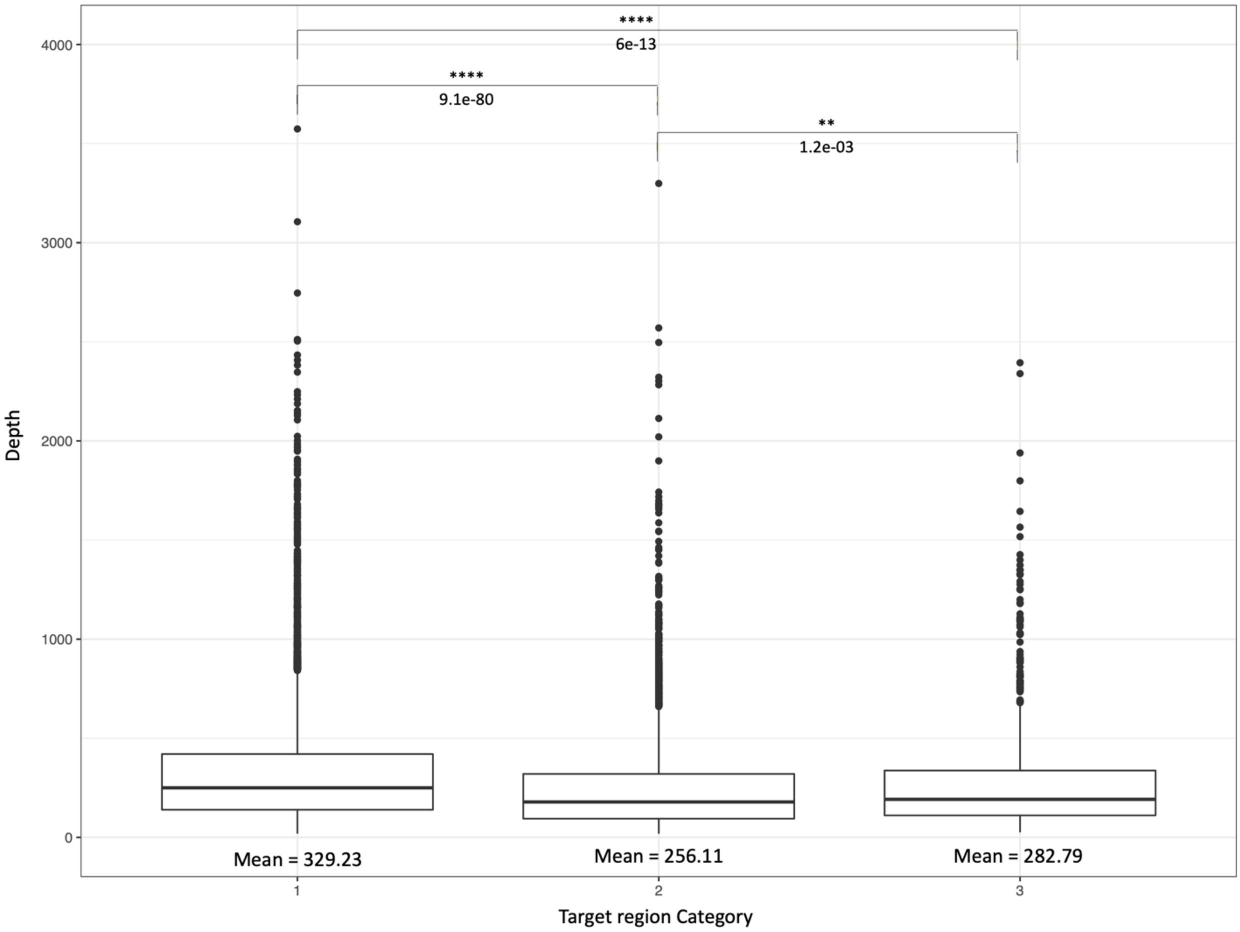


**Supplemental Figure S10. Distribution of average raw read depth per probe set for the three target region categories.** Mean read depth of target regions across all 288 individuals is on the y-axis and the target region categories are on the x-axis. The significant values were determined using the Wilcoxon sign ranked test. ** $p < 0.01$ and **** $p < 0.0001$.

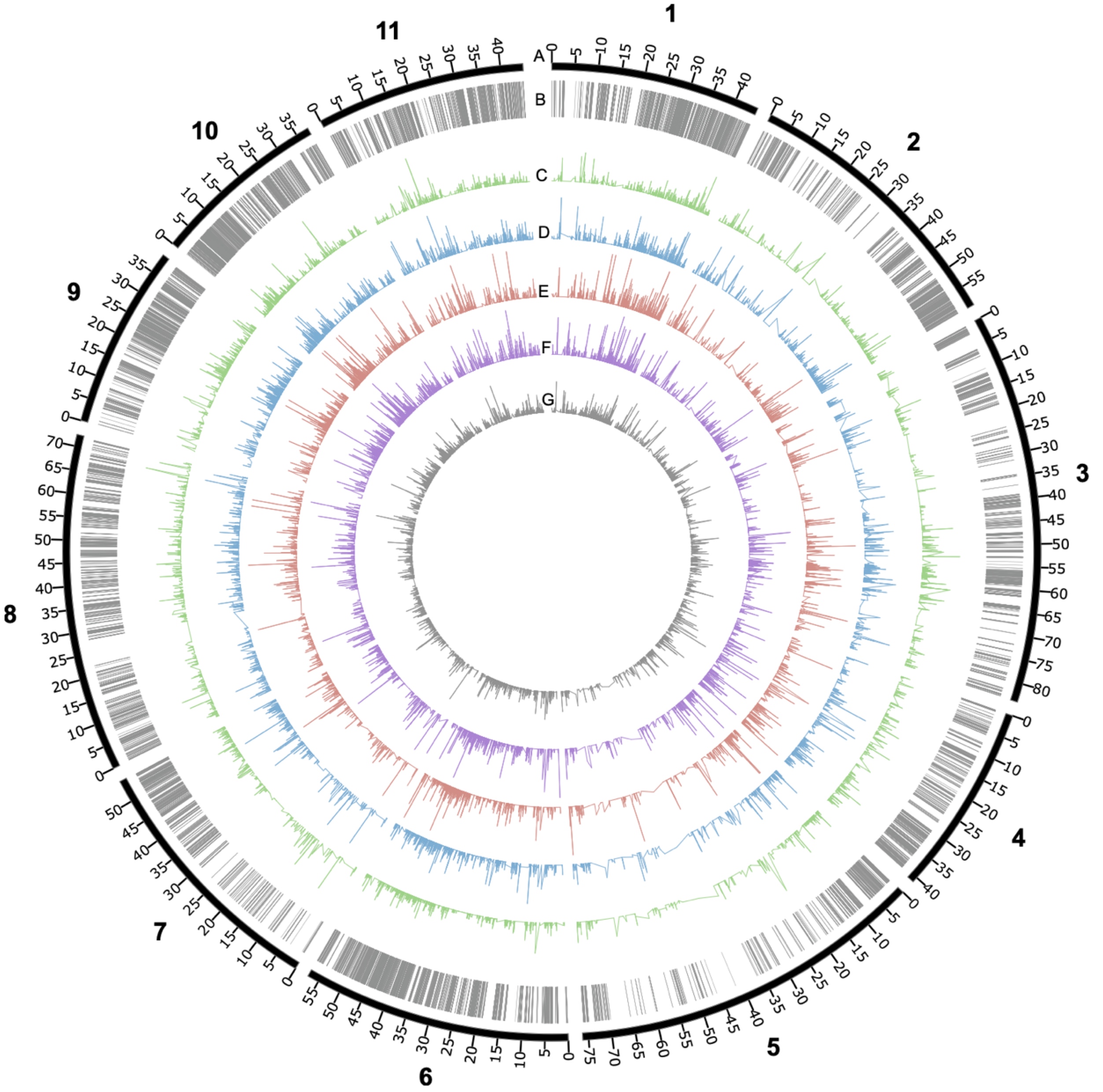


**Supplemental Figure S11. Percent of individuals with target regions containing three or four haplotypes per individual (tetraploid calls) in the four species. A.** Megabase values and chromosomes based on the *E. grandis* v2 reference genome (Bartholomé et al., 2015). **B.** Physical position of the target regions. **C.** Percent of target regions containing three or four haplotypes across 20 *E. grandis* individuals (range 0% - 100%). **D.** Percent of target regions containing three or four haplotypes across 20 *E. urophylla* individuals (range 0% - 100%). **E.** Percent of target regions containing three or four haplotypes across 20 *E. dunnii* individuals (range 0% - 100%). **F.** Percent of target regions containing three or four haplotypes across 20 *E. nitens* individuals (range 0% - 100%). **G.** Percent of target regions containing three or four haplotypes across all 288 samples (range 0% - 84.73%).

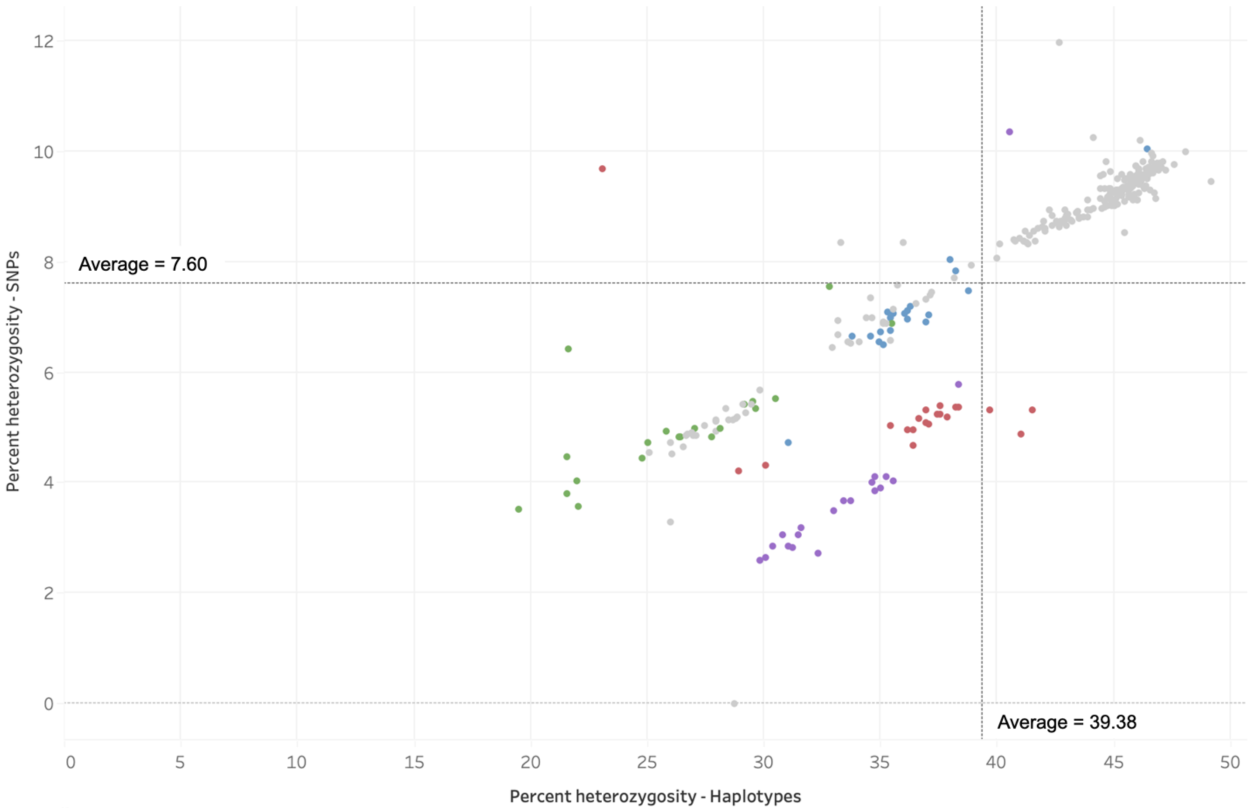


**Supplemental Figure S12. Average heterozygosity of individuals based on SNPs and haplotypes in Category 1 target regions.** The percent of heterozygosity of SNPs (y-axis) and haplotypes (x-axis) for 288 individuals. Each dot represents an individual. The average percent heterozygosity for SNPs (horizontal line) and haplotypes (vertical line) is shown. Dots are coloured according to species, *E. dunnii* red, *E. grandis* green, *E. urophylla* blue, *E. nitens* purple and the remaining individuals in grey are full-sib progeny and parents. Prior to calculating haplotype heterozygosity, target regions in individuals with two or more haplotypes were marked as missing.

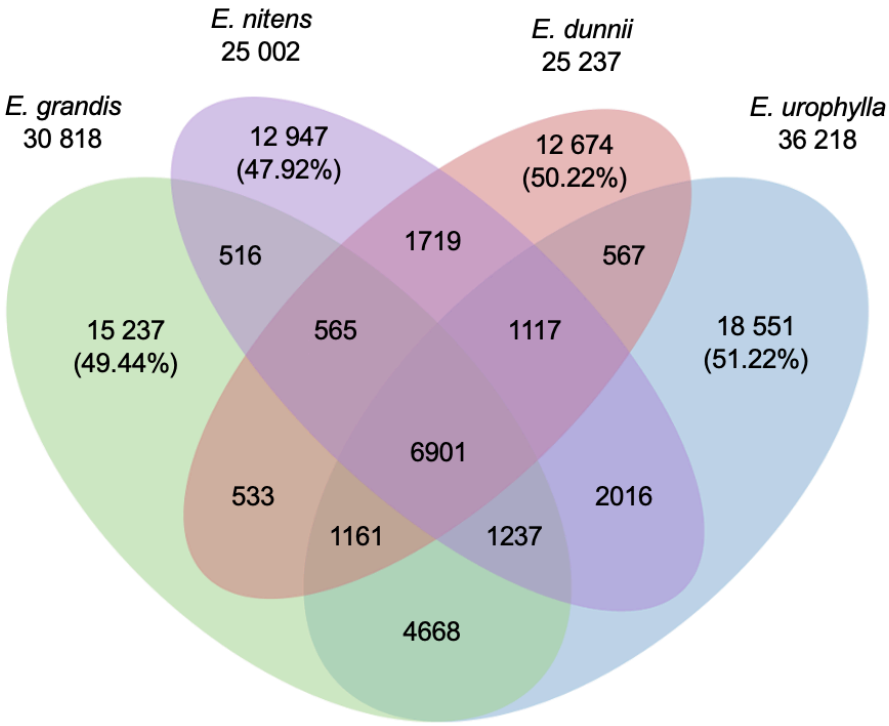


**Supplemental Figure S13. Number of shared and unique high quality haplotypes from Category 1 target regions in the four species.** The numbers underneath the species names are the number of haplotypes present in the species. The percent of unique haplotypes per species is shown. Prior to the analysis, individual haplotype calls that contain three or four haplotypes were converted to missing.

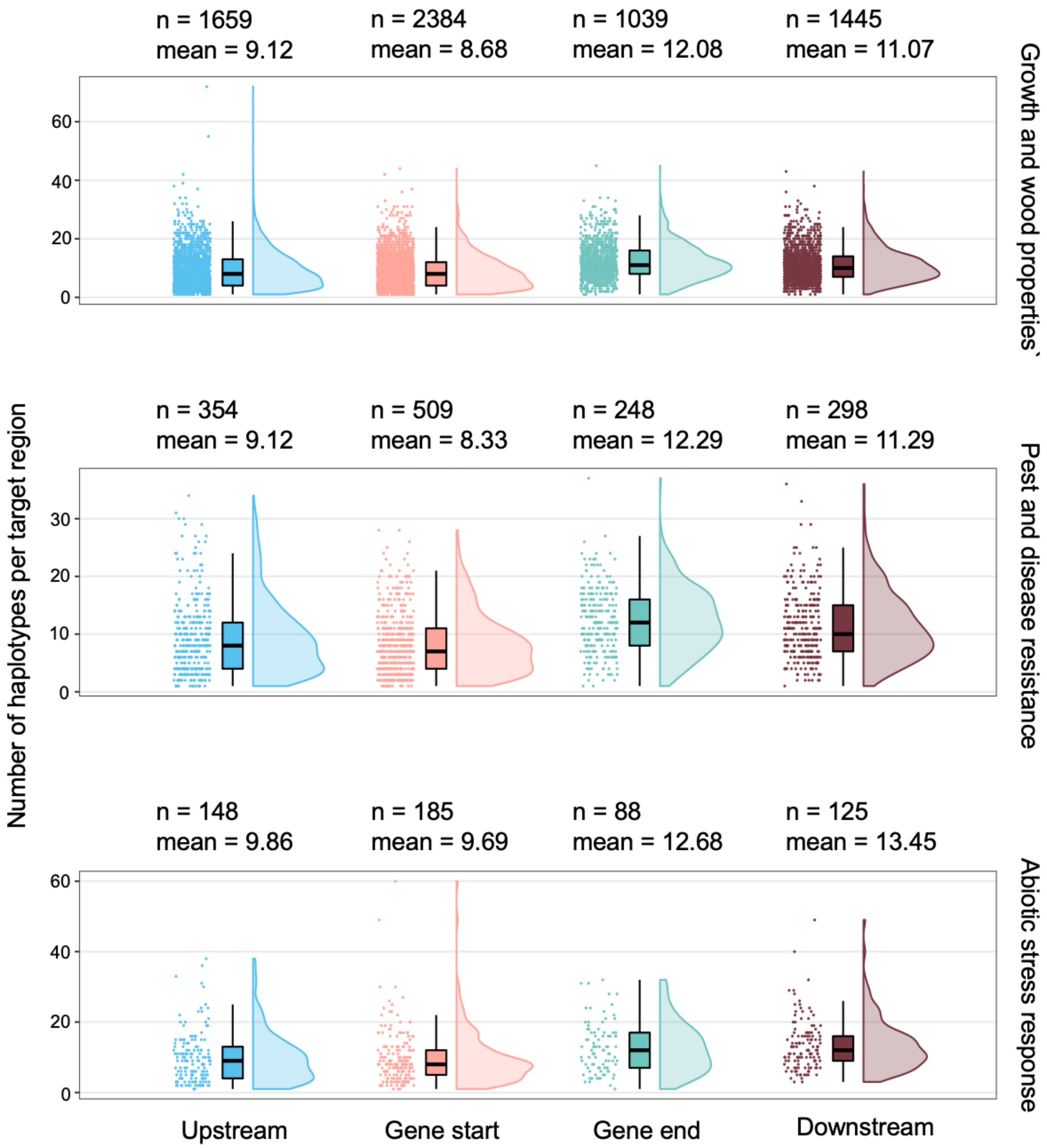


**Supplemental Figure S14. Haplotype diversity across gene categories and gene regions**. The number of haplotypes per target region (y-axis) was recorded for each of the four gene regions (x-axis). A total of 80 individuals (four species) were analysed per target region making the theoretical maximum haplotypes equal to 160 per target region. The mean value shown above each graph is the average number of haplotypes per target region and n is the number of target regions analysed in each category

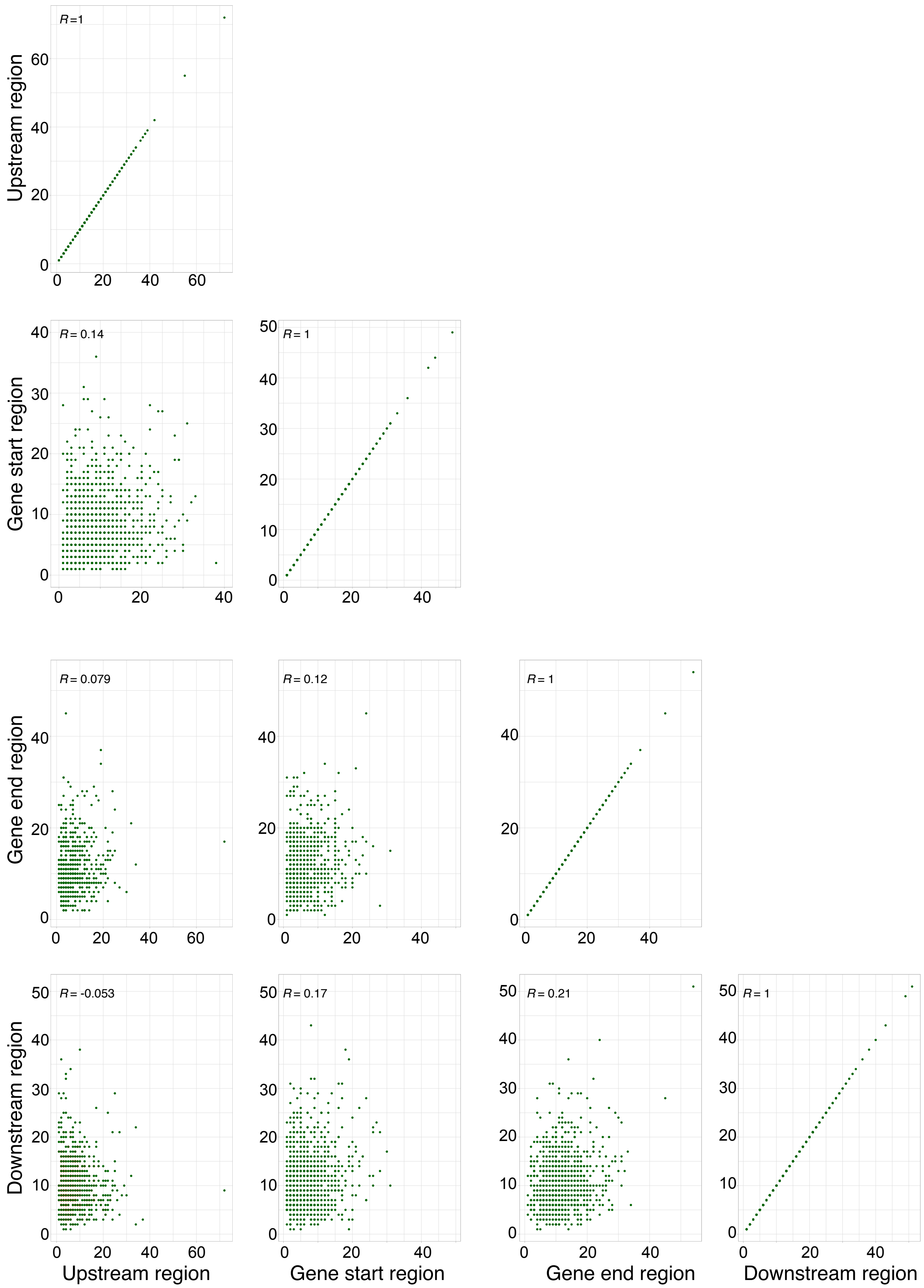


**Supplemental Figure S15. Pairwise correlations of haplotype numbers (proxy for haplotype diversity) among the four gene regions.** Each dot represents a Category 1 target region.

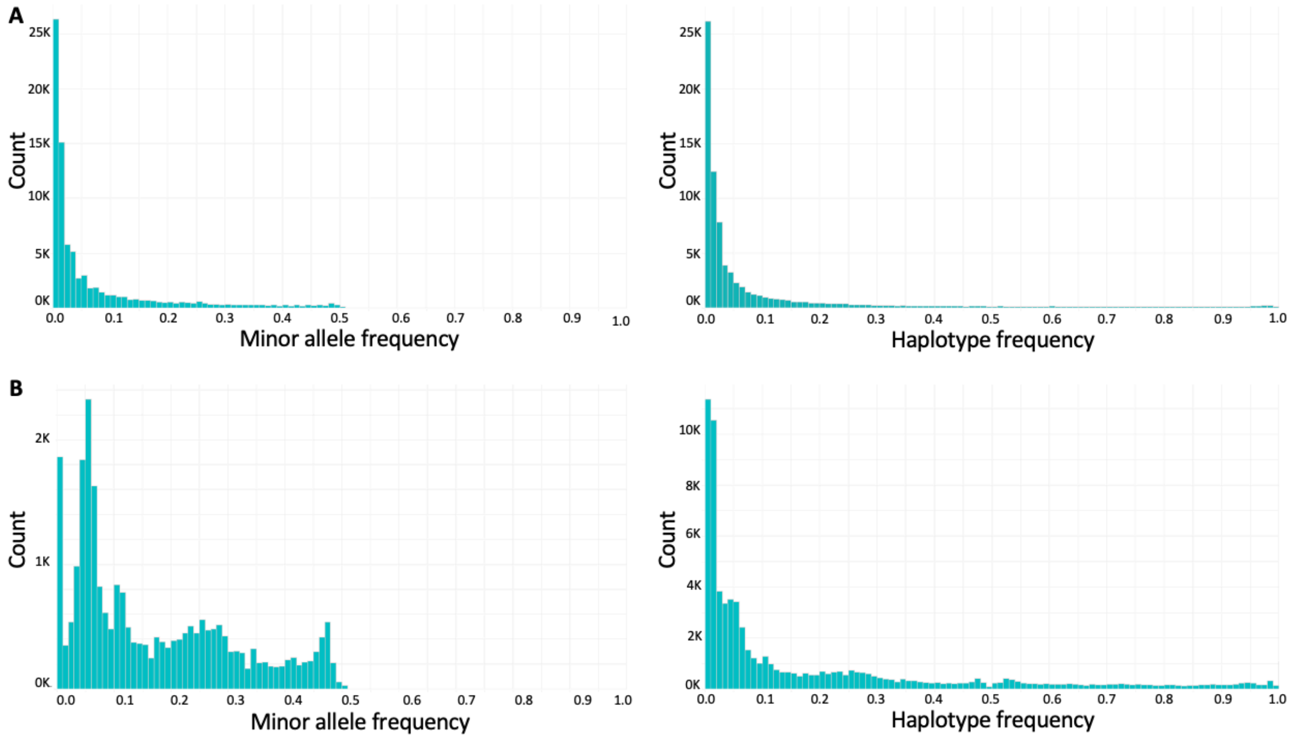


**Supplementary Figure S16. Distribution of allele frequency for haplotypes and SNPs. A.** SNP minor allele frequency (left) and haplotype frequency (right) for Category 1 target regions in all four species. **B.** SNPs minor allele frequency (left) and haplotype frequency (right) for Category 1 target regions in a HS family.

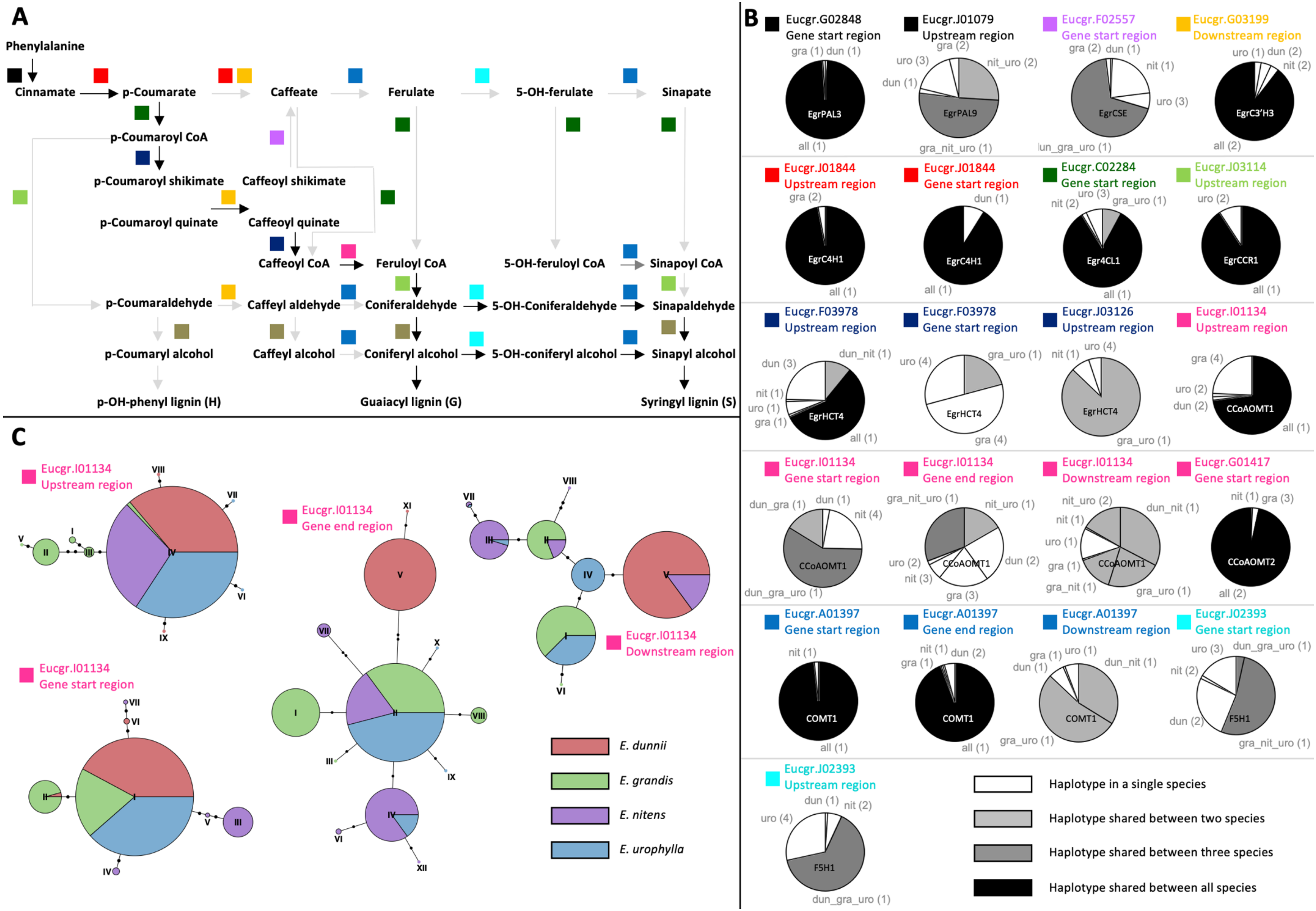
A
Phenylalanine
Cinnamate
p-Coumarate
Caffeate
Ferulate
5-OH-ferulate
Sinapate
p-Coumaroyl CoA
p-Coumaroyl shikimate
Caffeoyl shikimate
p-Coumaroyl quinate
Caffeoyl quinate
Caffeoyl CoA
Feruloyl CoA
5-OH-feruloyl CoA
Sinapoyl CoA
p-Coumaraldehyde
Caffeyl aldehyde
Coniferaldehyde
5-OH-Coniferaldehyde
Sinapaldehyde
p-Coumaryl alcohol
Caffeyl alcohol
Coniferyl alcohol
5-OH-coniferyl alcohol
Sinapyl alcohol
p-OH-phenyl lignin (H)
Guaiacyl lignin (G)
Syringyl lignin (S)
B
Eucgr.G02848 Gene start region
EgrPAL3
Eucgr.J01079 Upstream region
EgrPAL9
Eucgr.F02557 Gene start region
EgrCSE
Eucgr.G03199 Downstream region
EgrC3'H3
Eucgr.J01844 Upstream region
EgrC4H1
Eucgr.J01844 Gene start region
EgrC4H1
Eucgr.C02284 Gene start region
Egr4CL1
Eucgr.J03114 Upstream region
EgrCCR1
Eucgr.F03978 Upstream region
EgrHCT4
Eucgr.F03978 Gene start region
EgrHCT4
Eucgr.J03126 Upstream region
EgrHCT4
Eucgr.I01134 Upstream region
CCoAOMT1
Eucgr.I01134 Gene start region
CCoAOMT1
Eucgr.I01134 Gene end region
CCoAOMT1
Eucgr.I01134 Downstream region
CCoAOMT1
Eucgr.G01417 Gene start region
CCoAOMT2
Eucgr.A01397 Gene start region
COMT1
Eucgr.A01397 Gene end region
COMT1
Eucgr.A01397 Downstream region
COMT1
Eucgr.J02393 Gene start region
F5H1
Eucgr.J02393 Upstream region
F5H1
Haplotype in a single species
Haplotype shared between two species
Haplotype shared between three species
Haplotype shared between all species
C
Eucgr.I01134 Upstream region
Eucgr.I01134 Gene end region
Eucgr.I01134 Downstream region
Eucgr.I01134 Gene start region
E. dunnii
E. grandis
E. nitens
E. urophylla

**Supplemental Figure S17. Haplotype diversity of genes in the lignin biosynthetic pathway. A.** Lignin biosynthetic pathway adapted from Carocha *et al.* (2015) with coloured squares representing unique protein functions using the same colours as in Carocha *et al.* (2015). **B.** Rate of haplotype sharing across species. Shown in each pie is the number of individuals carrying haplotypes shared or unique across species. White represents haplotypes present in a single species, light grey represents haplotypes shared between two species, dark grey represents haplotypes shared between three species and black represents haplotypes shared across all species. The species are denoted as *E. grandis* (gra), *E. urophylla* (uro), *E. dunnii* (dun) and *E. nitens* (nit) with combinations of species represented with an underscore. The numbers in brackets represent the number of haplotypes. The text inside the pie charts represent the gene name. **C.** Haplotype networks for Eucgr.I01134 for the four gene regions. Each colour represents a species, with the size of the section representing the number of individuals.
